# Structure-function studies of HRI_KD-ΔKI_, a Minimal Kinase Domain of Human Heme-Regulated Inhibitor Kinase

**DOI:** 10.64898/2026.07.06.735516

**Authors:** Mohan B Rajasekaran, Jessica Booth, Damien F. Crepin, S. Mark Roe, Lihong Zhou, Rohanah Hussain, Tiberiu-M Gianga, Giuliano Siligardi, Ramon Gonzalez-Mendez, Maria Staikopoulou, Haitham Hassan, Antony W. Oliver, Erika J. Mancini, John Spencer

## Abstract

EIF2α kinase heme-regulated inhibitor (HRI) is a novel target for haematological malignancies with modulators reported to trigger cell death *via* the HRI-eIF2α-ATF4 pathway. We report a protocol for producing the minimal kinase domain of full-length human HRI, termed ‘HRI_KD-ΔKI_,’ where the unstructured 140 amino acid (aa) kinase insert (KI) within HRI kinase domain (HRI_KD_) is replaced with a 2aa glycine/serine (GS) linker. X-ray crystal structures were determined of ‘apo’-HRI_KD-ΔKI_ and of its complex with ATP at 2.1 & 2.5 Å resolution respectively. Both structures display a canonical bi-lobal kinase fold. However, they remain in a non-productive state with a displaced C-helix, disassembled R-spine, and a disordered activation segment hindering the substrate site. Biophysical assays (fluorescence based thermal shift & Synchrotron Radiation Circular Dichroism) demonstrate HRI_KD-ΔKI_ retains its functional ligand-binding conformation. All together, these findings define structural and ligand-binding features of HRI to support ongoing drug discovery efforts in blood cancer.

## INTRODUCTION

Heme-regulated inhibitor kinase (HRI) is a key sensor of cellular stress and a member of the conserved family of eIF2α kinases that initiate the Integrated Stress Response (ISR).^1^ Originally characterised in rabbit reticulocytes as a heme sensor, HRI is activated during heme deficiency and phosphorylates Ser51 of the α-subunit of eukaryotic translation initiation factor 2 (eIF2α) yielding eIF2αP.^2,3^ This phosphorylation event acts as a switch for the ISR, which simultaneously inhibits globin protein synthesis, enhancing the translation of regulatory factors such as Activating Transcription Factor 4 (ATF4), through the HRI-eIF2α-ATF4 pathway.^4,5^ HRI is important in hemeoglobinopathies and erythropoiesis, coordinating globin protein synthesis and heme availability.^6,7^ HRI also controls the innate immune response^8^, mitochondrial stress response ^9–11^ and cytoplasmic and neuronal proteostasis.^12,13^

Cancer cells often have elevated iron/ferritin levels^14,15^ relying on heme associated proteins^16^ for proliferation and survival where protein synthesis ^17^ occurs *via* the ternary eIF2-GTP-Met-tRNAi (TC) complex. HRI is a therapeutic target in hematological malignancies because its activation aberrates TC complex formation, halting global protein synthesis, impacting cancer cell proliferation, and triggering apoptosis.^18^ HRI is expressed in sensitive and resistant multiple myeloma (MM1.S/.R) cells.^19^ Pharmacological activation of HRI by 1-(benzo[*d*][1,2,3]thiadiazol-6-yl)-3-(3,4-dichlorophenyl)urea, BTdCPU^20^, induced eIF2 phosphorylation and upregulated C/EBP-homologous protein (CHOP), promoting apoptosis in MM1.S and MM1.R cells. Similarly, HRI mediated eIF2α phosphorylation, either by (BTdCPU), or dihydroartemisinin (DHA) treatment and induced apoptosis in murine BCR-ABL^+^ B-ALL (B-precursor acute lymphoblastic leukaemia) and human Ph^+^ (Philadelphia positive) leukaemia cell lines.^21^ HRI operates via the HRI-eIF2αP-ATF4 pathway by repressing expression of the anti-apoptotic protein myeloid cell leukaemia-1 (MCL-1). Studies further demonstrated that HRI deficiency (by genetic ablation) in the above cell lines abrogated the apoptotic effect, underpinning the crucial role of HRI in the regulation of apoptosis in B-ALL. Moreover, HRI activation (with BTdCPU or DHA) synergised with Bcl-2 Homology 3 (BH3) mimetics, inducing apoptosis in mouse and human leukaemia cell lines as well as patient derived xenografts of human B-ALL (Ph+ and Ph-like).^21^

The importance of HRI in MDS-RS (Myelodysplastic syndrome with ring sideroblasts) patient samples was demonstrated where Splicing Factor 3b Subunit 1 (Sf3B1) mutation activated the HRI-induced response pathway with abnormal terminal erythroid differentiation.^22^ Deletion/downregulation of HRI led to ATF4 mediated upregulation of expression of genes involved in erythroid differentiation, potentially diminishing anaemia and transfusion burden for MDS-RS patients.

These observations have inspired studies validating HRI as a druggable target with the observation of potential therapeutics, including HRI activators, BTdCPU, cHAUS (1-phenyl-3-((1,4-*trans*)-4 phenoxycyclohexyl)urea)^23^, nucleoside mimetic compounds^24^ and inhibitors including aminopyrazolidanes ^25^, known kinase inhibitors like dabrafenib, encorafenib, bosutinib and GCN2iB^26,27^ as reference small-molecule ligands for HRI.

HRI belongs to the eIF2α kinase family,^28^ alongside PKR (protein kinase RNA-activated),^29^ GCN2 (general control non-derepressable 2),^30^ PERK (PKR-like ER kinase).^31^ All three related kinases have been structurally elucidated by X-ray crystallography, but the structure of HRI remains elusive.^32,33^ This represents a critical bottleneck for developing small molecule therapeutics, as a molecular roadmap of HRI’s features would be key in revealing its overall fold, the binding mode of ligands, and druggable allosteric hot spots. HRI is a 630 aa protein with a complex architecture including three subdomains (Figure 1), a ∼100 aa globin-like N-terminal (NTD), a ∼425 aa core kinase domain (KD), between aa residues 161 to 585, and ∼50 aa C-term coiled domain (CC). Additionally, it contains a unique ∼140 aa kinase insertion sequence (KI, 240-376 aa) within the kinase domain. HRI’s function is further regulated by heme-binding, through a stable ‘S’ site in HRI’s NTD domain and a reversible site ‘R’ in the KD domain. His119 and His120 (from NTD) and Cys411 (from KD) identified as potential key heme-binding motifs^34^ as well as two potential heme regulatory motifs (HRM), designated as HRM1 (residues 410-415) and HRM2 (residues 552-557),^3^ with a Cys-Pro (CP) motif in the C-lobe of kinase.

**Figure 1.**
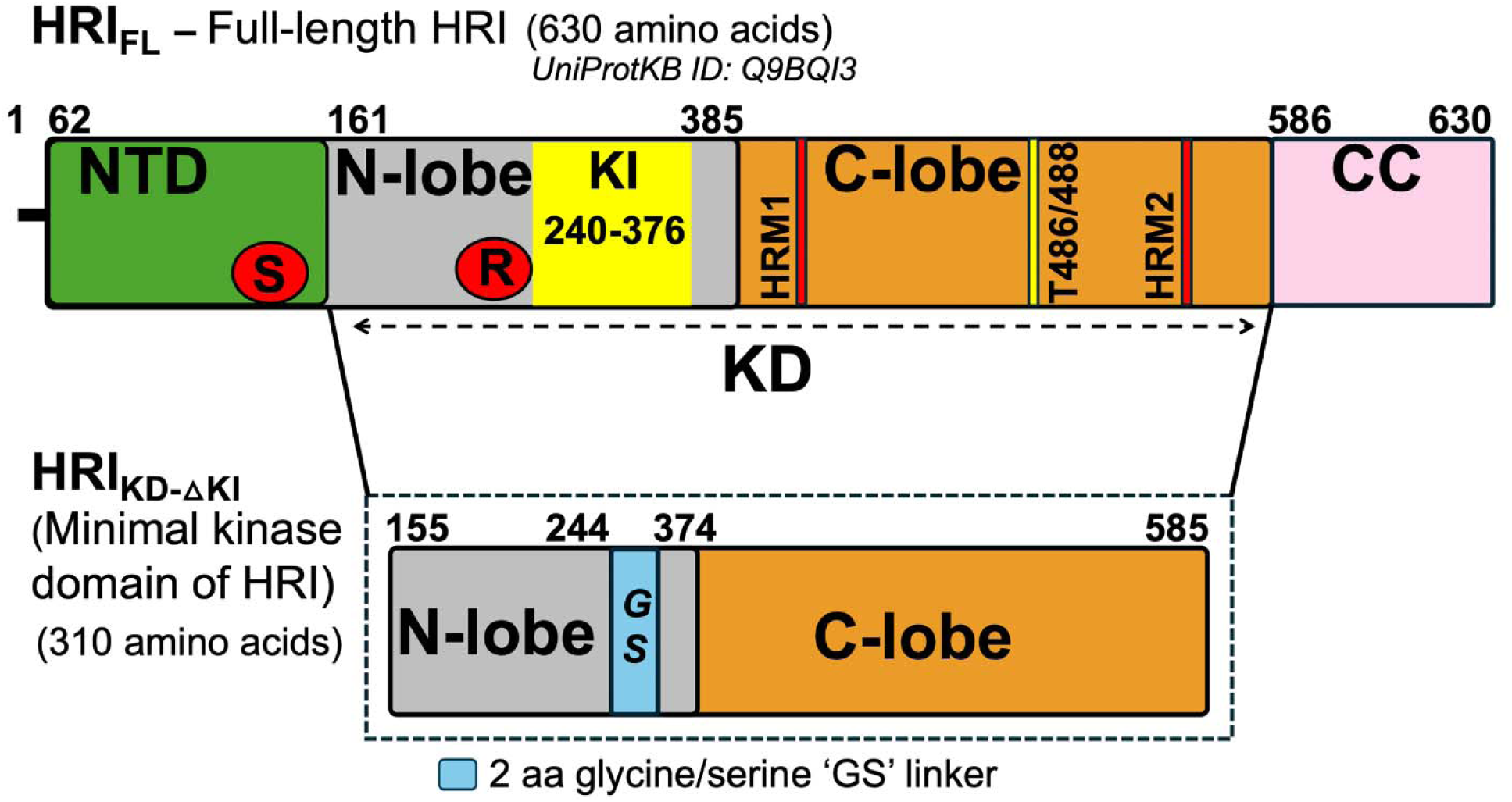
**Domain topology of Full-Length HRI (HRI_FL_) and Design of the HRI_KD-_**_Δ**KI**_. HRI comprises 630 aa and is structurally defined by three subdomains: the ∼100 aa N-term globin-like domain (NTD), the ∼425 aa core kinase domain (KD) comprising N- and C-lobes and the ∼50 aa C-term coiled domain (CC). The KD domain also contains a unique ∼140 aa kinase insertion sequence (KI, 240-376 aa) within the kinase domain. Key features include: two distinct heme binding sites (red circles) -a stable (S) site in the NTD and a reversible (R) site in the KD; the key activation loop residues in KD are, Thr486 and Thr488, (yellow thick line), which are auto phosphorylated; two unique heme regulatory motifs, HRM1 and HRM2 (red thick lines), in the C-lobe of KD. HRI_KD-ΔK**I**_ is a ∼35.3 kDa construct that models the minimal kinase domain by deleting the unstructured KI domain (245-373), replaced by a 2-amino acid glycine/serine (GS) linker. Domains not to scale.

Here, we disclose a robust protocol for the expression and purification of a minimal kinase domain of human full-length HRI (HRI_FL_), designated as HRI_KD-ΔKI_, which models the core kinase domain of HRI (HRI_KD_) (Figure 1), where the unstructured KI domain is replaced with a 2 aa glycine/serine (GS) linker, inspired by our previous protocols.^35,36^ Next, we will present biochemical and biophysical characterization of HRI_KD-ΔKI_ to offer valuable insight into its folding, stability, oligomeric state and provide crucial preliminary data on the ligand binding/affinity and ligand-induced conformational changes with known reference small- molecule ligands. Finally, we have solved the X-ray crystal structures of ‘*apo*’ HRI_KD-ΔKI_ (labelled hereafter as simply HRI_KD-ΔKI_) and its complex with ATP (adenosine-5′-triphosphate) to 2.1 and 2.5 Å resolution respectively. We also highlight a few salient structural markers observed in our structures which have c-Src^37^/cAbl^38^ like inactive-like conformation features. The structural framework of unique HRM motifs, the non-canonical conformation of G-helix, shared with other eIF2α kinase family members, is also discussed.

In summary, the structure-function studies presented here reveal the potential of HRI_KD-ΔKI_ as a reliable structural and functional platform to accelerate ongoing small molecule drug discovery efforts, particularly in blood cancer.

## RESULTS

### Design, expression and purification of HRI_KD-ΔKI_

We initially attempted to express HRI_FL_ as recombinant N-term GST/His-fusions either in *Escherichia coli (E. coli)* or *Spodoptera frugiperda* (*S. frugiperda)* cells and produced N-term His tagged HRI_FL_ from *E. coli* (∼80-90% purity) on a small scale (data not shown). However, both systems proved problematic, including poor expression profiles, batch-to-batch reproducibility, low protein yield, significant co-expression of non-specific proteins. These issues prevented high-resolution structural studies and biochemical/biophysical functional characterization of full-length HRI.

*In silico*-based analysis of the HRI_FL_ protein sequence (IntFOLD^39^, AlphaFold^40^, PrDOS^41^) predicted a kinase insert sequence of approximately 140 aa at positions 240 to 376 (Figure S1) within the KD domain, to be intrinsically disordered with a high Ser (17%) and Glu (14%) content. To enable the structure-function studies of HRI, next, we designed a minimal kinase domain of HRI, termed HRI_KD-ΔKI_, where a large segment of the KI domain, aa residues 245-373, was deleted and replaced with a two aa ‘GS’ linker (Figure 1). Using this construct, we were readily able to express in *E. coli* and purify the recombinant protein in milligram quantities by means of two-step purification using standard chromatographic techniques. (Figure S2A). The identity of the HRI_KD-ΔKI_ was confirmed via mass spectrometry (MS) analysis of a tryptic gel band digest of HRI_KD-ΔKI_ (BBSRC Mass Spectrometry and proteomics Facility, University of St. Andrews, UK). Second, deconvolution of the representative mass spectrum from in-house denaturing intact LC-MS studies on HRI_KD-ΔKI_ revealed a dominant peak at 35.7 kDa with 100% relative abundance (Figure S2D), which closely matches with expected theoretical MW of 35.3 kDa for HRI_KD-ΔKI_.

### Biochemical and Biophysical characterisation of HRI_KD-ΔKI_

The oligomeric state of HRI_KD-ΔKI_ was initially assessed by size-exclusion chromatography (SEC). The calibrated SEC experiments confirmed HRI_KD-ΔKI_ to be predominantly monomeric in solution and its elution (Figure S2B) corresponded to relative molecular weight (MW) of 36.3 kDa consistent with its expected theoretical MW of 35.3 kDa (Figure S2B and S2C).

Next, we examined the phosphorylation status of our bacterially expressed HRI_KD-ΔKI_ by mass spectrometry (Proteomics Facility, University of Bristol, UK). The peptides from protease digests revealed >20 modified phospho-sites (Table S1). Out of 27 sites, Tyr 193, Thr486, Thr488, Thr493 were a few of the key residues among them that have been reported as critical for autophosphorylation activity/eIF2alpha kinase activity based on site-directed mutagenesis studies.^42^ These residues were also identified in the previous MS studies of HRI_FL_ of mouse/human HRI.^26,42,43^ Interestingly, phosphorylation at Ser409 was another observed modification and was subsequently resolved in our X-ray crystal structure (*vide infra*). In addition to the above primary sites mapped within the native sequence of HRI_KD-ΔKI_, two to five supplementary sites also mapped onto the remnant of the HRV-3C cleavage site and engineered linker sequence respectively.

We next measured the kinase activity of HRI_KD-ΔKI_ and compared it with full-length HRI, HRI_FL-RB_, using a radiometric assay with casein as model substrate,^44,45^ with velocities plotted against ATP concentration (Figure S3). HRI_FL-RB_ exhibited higher kinase activity than the HRI_KD-ΔKI_ across the ATP concentration range tested, while the HRI_KD-ΔKI_ retained measurable residual activity.

In order to shed more light on the folded state and secondary/tertiary features, we subjected purified HRI_KD-ΔKI_ protein to a Fluorescence based thermal shift assay (FTS) and SRCD (Synchrotron Radiation Circular Dichroism) assay. FTS assays with HRI_KD-ΔKI_ in its *apo*-form showed a single transition (Figure 2A) during thermal unfolding with a mean T_m_ of 58.0 °C compared to ∼50 CC reported for *apo* HRI_FL_.^26,46^ The assay also confirmed that an ATP-binding pocket of HRI_KD-ΔKI_ is accessible, with ATP and AMP-PNP stabilising HRI_KD-ΔKI_ with a ΔT_m_ > +1.0 °C (Figure 2A, Table 1).

**Figure 2:**
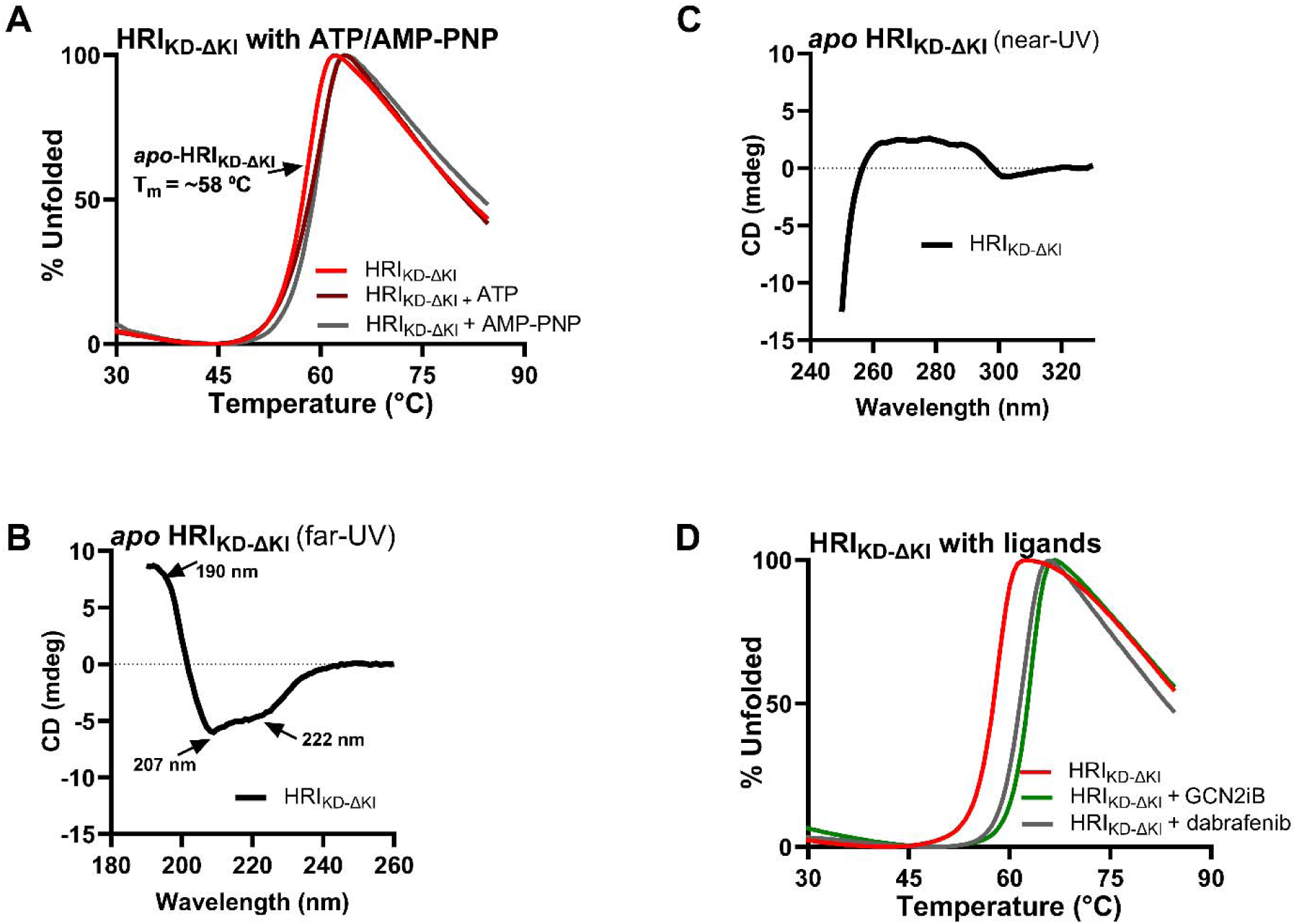
**Biophysical validation of secondary/tertiary fold and characterization of ATP binding pocket accessibility of HRI_KD-_**_Δ**KI**_ (A) Representative FTS based normalized thermal denaturation profile for HRI_KD-ΔKI_ (red) (single-phase transition denaturation profile with a T_m_ around ∼58.0 CC and shift upon binding the substrate ATP (brown) and AMP-PNP (grey), indicating domain stabilization (ΔT_m_ of > +1CC, Table 1). (B) Far-UV SRCD spectra for the HRI_KD-ΔKI_ clearly displayed three distinct spectral bands characteristic of a well-folded, alpha-helix-rich conformation. (C) SRCD near-UV spectra of the HRI_KD-ΔKI_ probing the tertiary structure feature aspects of protein. (D) Representative thermal denaturation profile for HRI_KD-ΔKI_ (red) upon binding of GCN2iB (green) and dabrafenib (grey) showing binding-induced ΔT_m_ of +4-5 CC by FTS assay (Table 1). The spectrum shown in all four cases represents the final averaged data from three or four consecutive scans (n=3 or 4 scans)

**Table 1:**
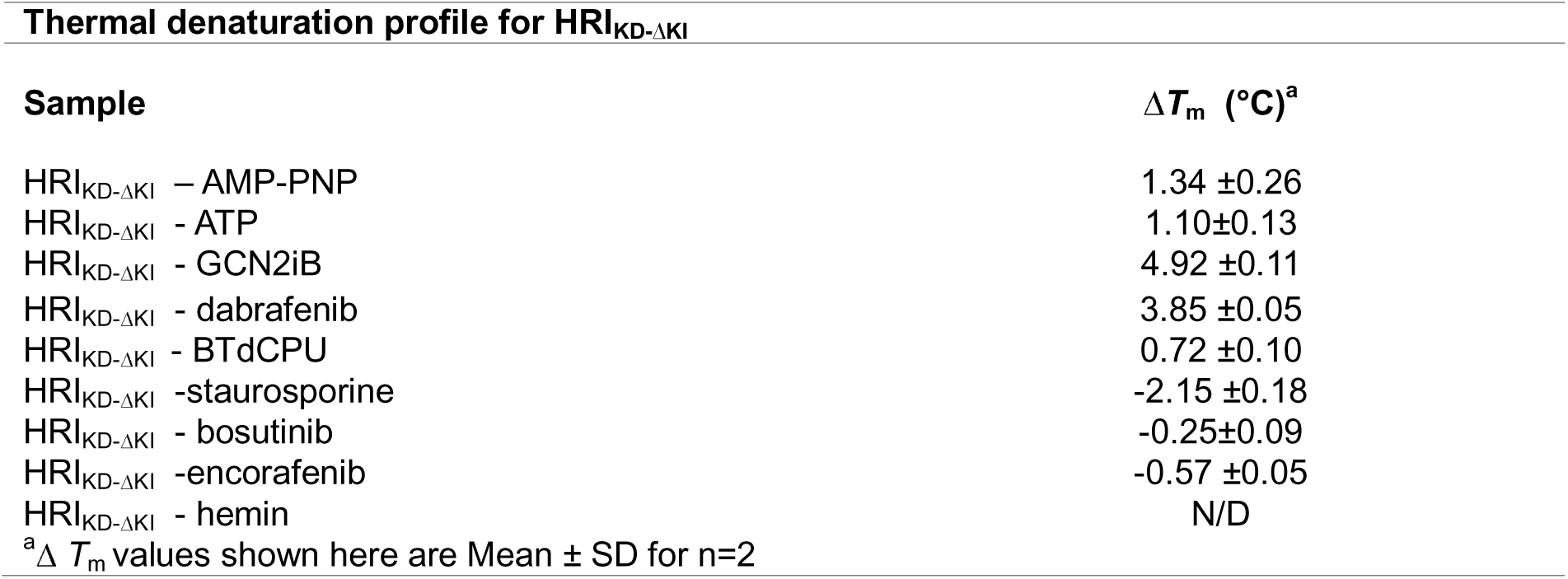
HRI_KD-ΔKI_ – substrate/ligand binding thermal denaturation studies by FTS assay.

| Thermal denaturation profile for HRI<sub>KD-ΔKI</sub> |  |
| --- | --- |
| Sample | $\Delta T_m$ (°C)<sup>a</sup> |
| HRI<sub>KD-ΔKI</sub> – AMP-PNP | 1.34 ±0.26 |
| HRI<sub>KD-ΔKI</sub> - ATP | 1.10±0.13 |
| HRI<sub>KD-ΔKI</sub> - GCN2iB | 4.92 ±0.11 |
| HRI<sub>KD-ΔKI</sub> - dabrafenib | 3.85 ±0.05 |
| HRI<sub>KD-ΔKI</sub> - BTdCPU | 0.72 ±0.10 |
| HRI<sub>KD-ΔKI</sub> -staurosporine | -2.15 ±0.18 |
| HRI<sub>KD-ΔKI</sub> - bosutinib | -0.25±0.09 |
| HRI<sub>KD-ΔKI</sub> -encorafenib | -0.57 ±0.05 |
| HRI<sub>KD-ΔKI</sub> - hemin | N/D |
| <sup>a</sup>Δ *T<sub>m</sub>* values shown here are Mean ± SD for n=2 |  |

Far UV SRCD spectra of HRI_KD–ΔKI_ display canonical α/β secondary structure features, with a positive band near ∼190 nm and negative minima at ∼207 and 222 nm (Figure 2B). Quantitative analysis indicated approximately 33% α-helix and 19% β-strand content, consistent with a mixed α+β structural fold (Figure S6). The near-UV SRCD (250-330 nm) spectra of HRI_KD-ΔKI_ revealed a broad peak between 260-300 nm (Figure 2C) due to contributions from the aromatic side chains (tyrosine, tryptophan and phenylalanine), confirming a stable tertiary structural fold. Taken together, these findings indicate the presence of well-defined secondary and tertiary structural features, confirming that HRI_KD-ΔKI_ adopts a properly folded state, and further establish the potential of biophysical assays (FTS/SRCD) for probing HRI_KD-ΔKI_ –ligand interactions using reference small-molecule ligands.

We next tested the binding of seven reference small-molecule ligands: BTdCPU, staurosporine, encorafenib, bosutinib, dabrafenib, GCN2iB and hemin, a chloride coordinated Fe(III) heme analogue (HRI inhibitor ligand) to HRI_KD-ΔKI_ by the FTS assay. Out of seven, GCN2iB, dabrafenib and BTdCPU thermally stabilised HRI_KD-ΔKI_ with positive ΔT_m_ values of approximately 5, 4 and 0.8 °C respectively (Table 1, Figure 2D & S4). An interesting observation was that dabrafenib and GCN2iB, with higher ΔT_m_ observed in our study, echoes FTS- based binding profiles reported for HRI_FL_ (ΔT_m_ of + 6.1 and + 9.1 °C respectively).^26^ However, non-binding or negative T_m_ shift binding was observed in common for the remaining four ligands (staurosporine, encorafenib, bosutinib and hemin) (Figure S4).

We further corroborated the HRI_KD-ΔKI_ – reference small-molecule ligand binding profile by far and near-UV SRCD assay. GCN2iB & dabrafenib (two top-most binding ligands from the FTS assay), hemin (HRI inhibitor ligand) and BTdCPU (HRI activator tool compound) were included in this assay. Ligand-dependent SRCD based thermal denaturation profiles reveal stabilisation trends (Figure S5). BTdCPU exhibits the elevated ellipticity (Figure S5A), which retained across the full temperature range (60–90 °C), whereas GCN2iB and dabrafenib produced moderate–strong stabilisation with positive ΔT_m_ values ∼2CC (Figure S5B). In contrast, hemin (Figure S5A) produced minimal effects, consistent with weak structural perturbation of the protein.

Ligand binding studies with far-UV SRCD induced modest yet reproducible spectral changes (Figure S5C and S5D), with BTdCPU increasing α-helical content and reducing β-strand contributions (Figure S6), whereas far-UV SRCD for hemin produced negligible perturbation (Figure S5C). These findings indicate ligand-induced local conformational adjustments without global unfolding, consistent with the maintenance of tertiary structural integrity. Near UV SRCD measurements of HRI_KD-ΔKI_ with small-molecule ligands revealed spectral changes between 300 to 370 nm confirming ligand interactions at the tertiary structural level (Figures S5E and S5F). Titration experiments for GCN2iB and dabrafenib generated clear induced CD signals at 338 and 355 nm respectively, enabling quantitative affinity determinations (Figure 3A-3B). The fitting of the data yielded apparent dissociation constants of ∼12.9 µM for GCN2iB and ∼7.6 µM for dabrafenib (Figure 3C-3D), indicating moderate binding affinity, which were reproducible across independent datasets confirming the robustness of the SRCD-based approach.

**Figure 3:**
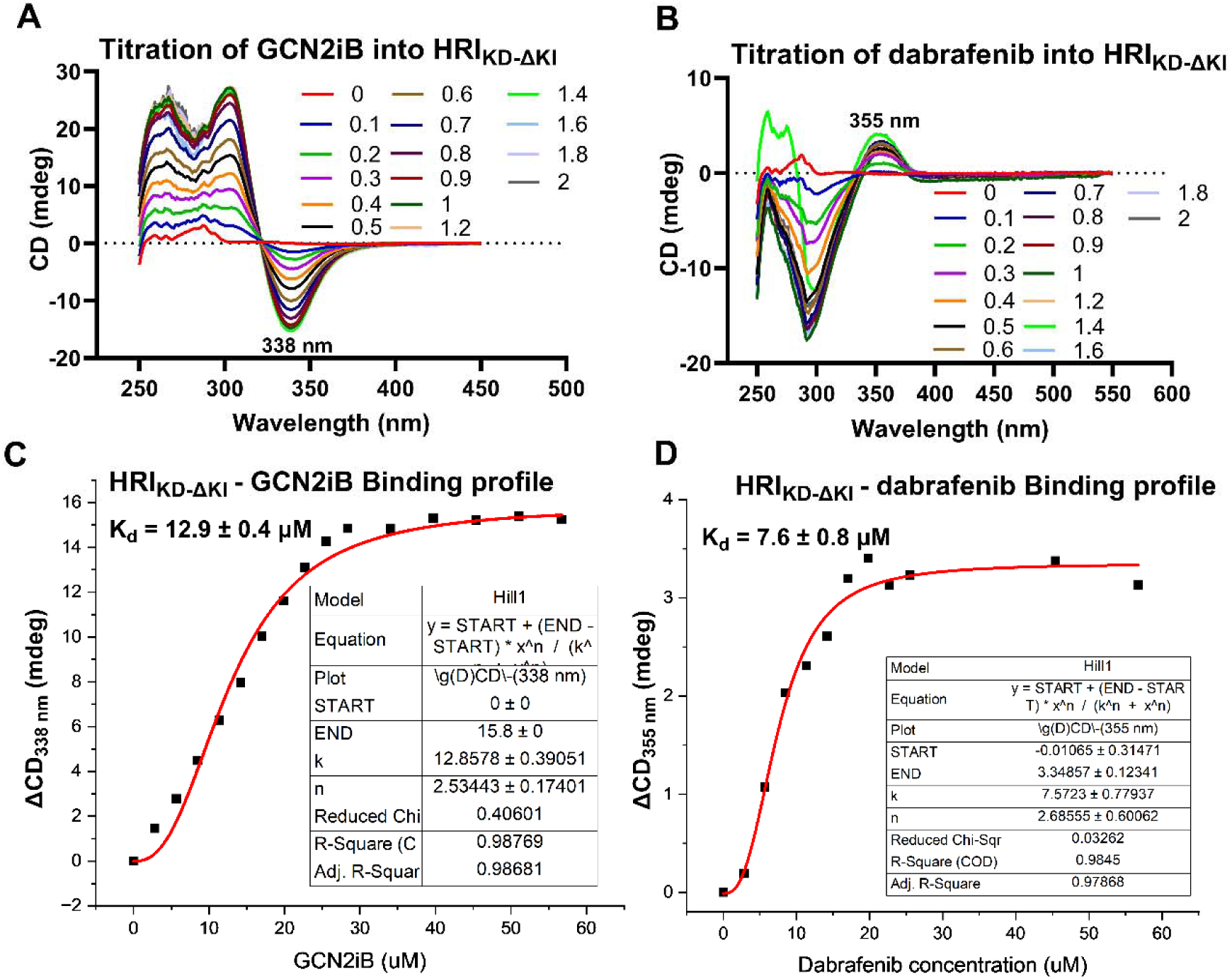
**Near-UV SRCD spectra for the HRI_KD-_**_Δ**KI**_ **– ligand titration studies and measurement of binding affinities of ligands by SRCD** (A-B) SRCD spectra of the titration of HRI_KD-ΔKI_ with GCN2iB and dabrafenib at different molar ratios respectively. Both ligands showed induced CD signals following the stoichiometric addition indicative of HRI_KD-ΔKI_ - ligand complex formation. The spectrum shown here represents the final averaged data from four consecutive scans (n=4 scans). (C-D) Binding curves were obtained by non-linear regression of mean CD signals at 338 nm (GCN2iB) and 355 nm (dabrafenib) across ligand titrations using Origin and GraphPad. The mean CD scan in each case was dilution-corrected and baseline-subtracted against buffer controls prior to fitting. Dissociation constants (K_d_) of 12.9 µM for GCN2iB and 7.6 µM for dabrafenib were determined using the Hill equation.

Integration of thermal, secondary, and tertiary structural datasets defines a consistent ligand interaction hierarchy. Dabrafenib and GCN2iB emerge as the strongest binders, supported by concordant affinity, CD perturbation, and stabilisation data. Collectively, these findings demonstrate that SRCD provides an integrated platform for probing binding affinity, conformational dynamics, and thermodynamic stabilisation in HRI_KD–ΔKI_ small-molecule ligand systems (Figure 2, 3 and S5).

### Crystallisation and structure determination of HRI_KD-ΔKI_ and HRI_KD-ΔKI_ -MgATP

HRI_KD-ΔKI_ crystals were obtained after three weeks incubation at 4 CC, diffracting to ∼2.1 Å resolution. HRI_KD-ΔKI_ – MgATP crystals were prepared by soaking *apo* HRI_KD-ΔKI_ with 5 mM MgATP and successfully diffracted in the resolution of 2.5 Å. The HRI_KD-ΔKI_ structure (with R_work_/R_free_ = 20%/23%, ∼2.1Å) was solved by molecular replacement with the GCN2 crystal structure (PDB code:1ZYC)^47^ as the search model. The details of crystallisation and structure determination are summarised in Methods & Table S2. The coordinates of HRI_KD-ΔKI_ were used as the starting model during refinement and modelling for the HRI_KD-ΔKI_ -MgATP structure elucidation.

HRI_KD-ΔKI_ and HRI_KD-ΔKI_-MgATP crystallised in the monoclinic space group P21 with unit cell dimensions; a = 56.37 Å, b = 78.14 Å, c = 90.17 Å, and α = 90C, β = 99.23C, γ = 90C & a = 56.85 Å, b = 80.08 Å, c = 88.19 Å, and α = 90C, β = 99.53C, γ = 90C respectively. The crystal asymmetric unit of both HRI_KD-ΔKI_ and HRI_KD-ΔKI_–MgATP structures contains two monomers (chains A and B), which are virtually identical, with low Cα RMSD values of 0.18 and 0.26 A° reported between chains A & B of each structure from pair-wise structure superimposition.

In this study, Chain A of the HRI_KD-ΔKI_ structure was investigated for detailed analysis of the bilobed kinase architecture, conserved motifs such as HRM, c-Src like inactive conformation whereas MgATP interactions were characterized using chain B of the HRI_KD-ΔKI_ –MgATP crystal structure complex. The structure-based superimposition at Cα atoms for both *apo* (chain A) and MgATP bound form (chain B) showed a rms deviation of ∼0.3 A°, indicating both forms adopt a similar conformation.

### Overall structural architecture of HRI_KD-ΔKI_ and HRI_KD-ΔKI_ -MgATP

The crystal structures of HRI_KD-ΔKI_ & HRI_KD-ΔKI_ -MgATP share a canonical kinase architecture with a bi-lobe kinase fold, possessing a small N-term lobe (N-lobe) and large C-term lobe (C-lobe) connected by a hinge region/loop with an ATP binding cleft positioned in between the lobes (Figure 4A).

**Figure 4:**
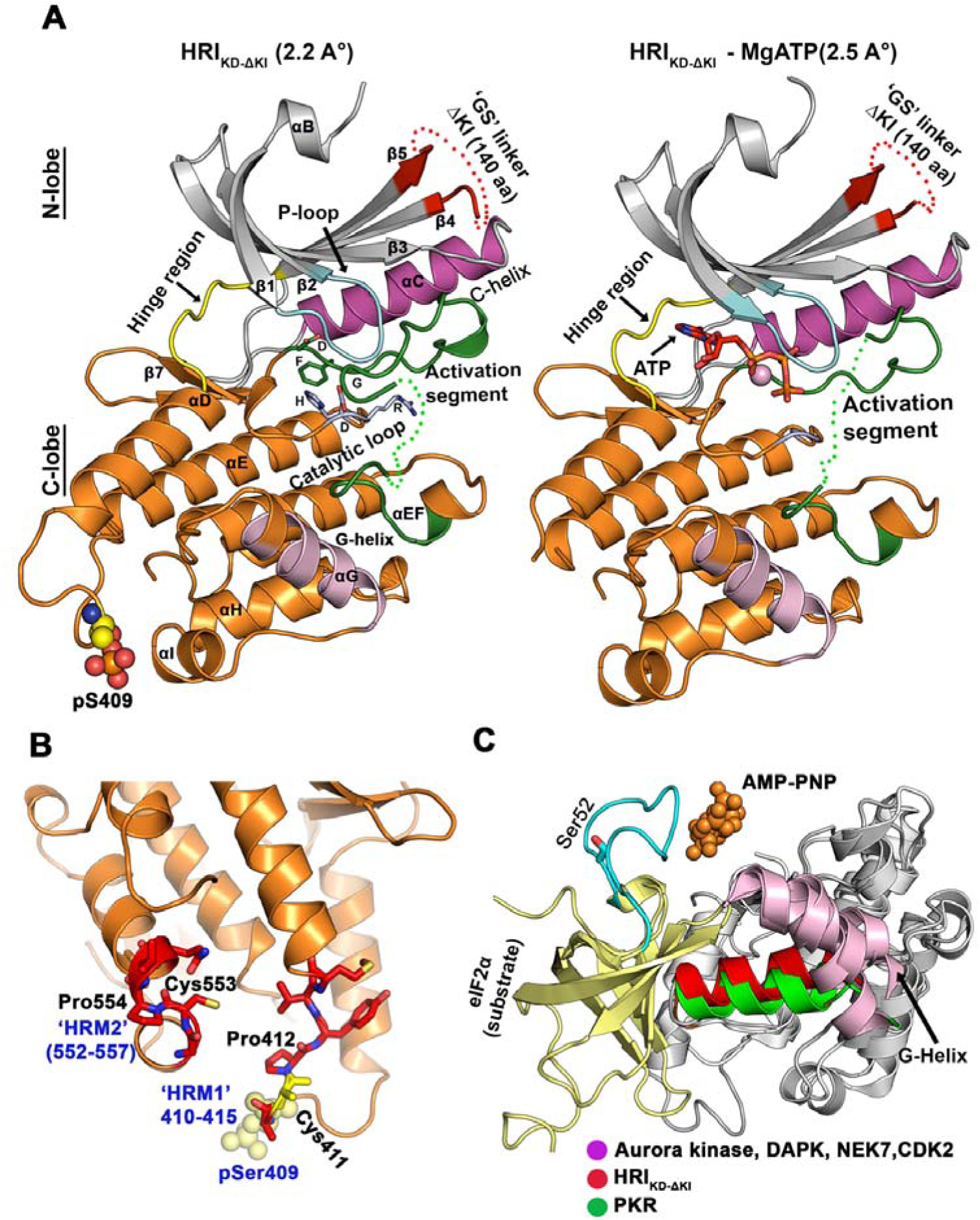
**Crystal structures for the HRI_KD-_**_Δ**KI**_ **and HRI_KD-_**_Δ**KI**_ **-MgATP forms.** (A) Cartoon representation of crystal structure of HRI_KD-ΔKI_ from *Homo sapiens*. HRI_KD-ΔKI_ displays a canonical bi-lobe kinase architecture with a small N-lobe (grey) and a large C-lobe (orange). The 2 aa GS linker (red) and part of the activation segment (green) are not modelled here and shown as dotted lines. The phosphorylated residue, pSer409 is shown as spheres. The key structural motifs such as ‘DFG’ and ‘HRD’ are well-defined in both structures and displayed in green and grey sticks. The crystal structure for HRI_KD-ΔKI_ - MgATP clearly reveals ATP (shown in orange sticks) between the N and C lobes. (B) Two potential heme regulatory motifs (HRM) are proposed for HRI located in the C-lobe. Both motifs possess a conserved, well-ordered, cysteine/proline motif (CP) and are labelled as HRM1 (aa residues 410-415) and HRM2 (aa residues 552-557). The thiol side chain for Cys 411 is not well-defined in our structure. The phosphorylated residue, pSer409 is in the vicinity of this HRM1 motif and it is shown as yellow spheres. (C) Close-up view of G-helix conformations of HRI and its closely related structural homologues. With PKR-eIF2α as reference structure, a displaced G-helix is observed for eIF2α kinase family members PKR (green cartoon), HRI_KD-ΔKI_ (red cartoon) and Aurora kinase/DAPK, NEK7, CDK2 (pink cartoon). AMP-PNP (from PKR) shown as orange spheres. As the phosphor-acceptor site of EIF2S1 (*Saccharomyces cerevisiae*) in PKR-EIF2S1 complex (2A19) is disordered, EIF2S1 from an AlphaFold model (Alphafold ID: AF-P05198-F1; *Homo sapiens*) is used here for the discussion on Ser52 of EIF2S1 (cyan).

Our HRI_KD-ΔKI_ structure revealed an N-lobe region that extends between aa residues 160-237 and 376-385, composed of a five stranded β-sheet (β1-β5) coupled with an αC- helix (C-helix) (Figure 4A). The flexible glycine rich loop (P-loop) between β1 and β2 is well-ordered. The C-helix (aa residues 204-220) is also well-ordered in the structure. Residues Lys196 and Glu214 are conserved and well-ordered in this structure, although quite distant from one another and no salt bridge is observed (approximately 10 Å apart; Figure 6). The engineered ‘GS’ linker, connecting between β4 and β5 of the N-lobe, replacing the deleted KI domain segment (aa 245-373), is poorly defined in the electron density map.

The larger C-lobe, stretching between aa residues 391-578, comprises seven α-helices (αD-αJ) and two β strands (β6 – β7), and is connected to the N-lobe by the hinge region (aa residues 386-390). The C-lobe mainly contains a docking site for substrates/proteins, a catalytic loop and activation segment. The ‘activation segment’ (aa residues 461-500) starts with a N-term located Asp-Phe-Gly (DFG) motif (Figure S7), comprising three sub-segments, a magnesium binding loop, activation loop, P+1 loop, ending with an ‘SPE’ (Ser-Pro-Glu) motif. The highly conserved DFG motif, starting with D461, is well-ordered. The middle part of the activation segment is the ‘activation loop’ (aa residues 468-489). Although clear electron density for aa residues 468-478 was observed, aa residues 479 to 492 were disordered and not modelled. Moreover, no significant electron density was detected for phosphor sites, especially Thr486, 488 and 493, identified in human/murine HRI as critical for catalytic activity^26,42,43^. The aa residues 493-500 of the P+1 loop were modelled in this structure (Figure S7).

In addition, a few more interesting structural features were observed in the C-lobe of the crystal structures of HRI_KD-ΔKI_. The first one is the successful mapping of the two potential heme regulatory motifs, HRM1 and HRM2,^3^ with a Cys-Pro (CP) motif, in our crystal structure. Both motifs, depicted as ‘HRM1’ and ‘HRM2’ in this study (Figure 4B), were well-ordered in our structure. However, no heme molecule binding event was observed at either HRM motif.

Secondly, the G-helix in the C-lobe (aa residues 529-540) is well-defined in the HRI_KD-ΔKI_ structure and adopts a non-canonical conformation (Figure 4C). Structural studies on the PKR-eIF2α complex ^48^ postulate that this distinct helical topology may assist in the optimal positioning of the phosphor acceptor substrate (eIF2α) to the active site and is consistent across with other eIF2α kinase family members, differentiating them from close HRI structural homologues such as Aurora kinase,^49^ CDK2,^50^ NEK7,^51^ and DAPK1,^52^ (from a DALI structure homologues search).

Thirdly, a unique αD-αE loop, which stretches between aa 403-415, where Ser409 is phosphorylated, with clear electron density observed (Figure S8A) is also observed in the HRI_KD-ΔKI_ crystal structure.

### Structural overview of HRI_KD-ΔKI_ -MgATP

This is the first detailed structural insight into HRI-substrate recognition with the ATP binding site exploration. The 2.5 Å resolution X-ray crystal structure of HRI_KD-ΔKI_ -MgATP, obtained by a soaking method, was refined to final R_work/_R_free_ of 23 and 27%. One ATP molecule and a Mg^2+^ cation are present in both molecules in the asymmetric unit.

The structure clearly reveals prominent density for the ATP molecule at the ATP cleft between the N- and C- lobes (Figure S8B). As expected, and observed in other kinases, the adenine ring of ATP is enclosed in a hydrophobic pocket formed at a domain interface comprising amino acids ligands from both the N-and C- lobes including Leu173, Val181, Ala194, Val227, Met384, Leu386, Cys387 and Phe449. The N1 and N6 amino groups of ATP form hydrogen bonds with the main-chain carbonyl/amino group of Gln385 and Cys387 of the hinge region (Figure 5). Similarly, the main chain carbonyl oxygen of the amino acid ligand Arg446 establish a hydrogen bond with the O2’ oxygen group of the ribose moiety of ATP. The stabilisation of α-phosphate of ATP is coordinated by the interaction of oxygen atoms, O1A & O2A with side chain amino/carboxamide group of Arg446 and Asn447 and magnesium atom (Figure 5). The β-phosphate of ATP is stabilised through oxygen atoms, O1B and O2B which interacts with Tyr 178 (from P-loop), Lys196 (from β3 strand), Asp461 (from DFG loop), magnesium and water (Figure 5). The final γ-phosphate is also stabilised by coordinated interaction between oxygen atoms of phosphate, O1G, O2G,O3G with Asp442(part of HRD motif), Asn447 and Lys444.The conformation of the P-loop of HRI_KD-ΔKI_-MgATP is slightly raised or ‘open’ in comparison to the *apo* state to accommodate ATP (Figure S9).

**Figure 5.**
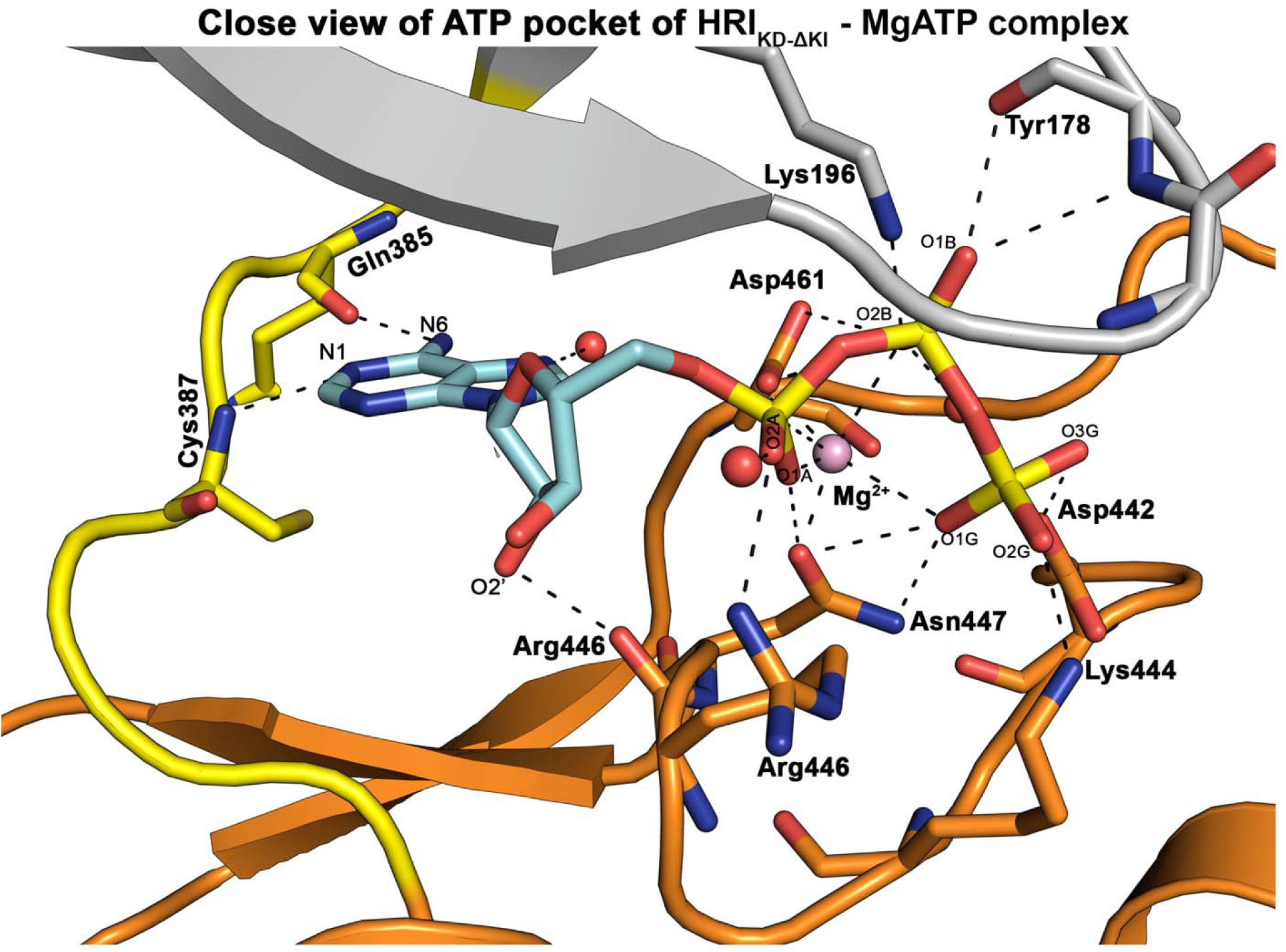
**Structural overview of the ATP binding pocket of HRI_KD-_**_Δ**KI.**_ The crystal structure of HRI_KD-ΔKI_ -MgATP was obtained by soaking ATP into apo-HRI_KD-ΔKI_ crystals using standard crystal soaking procedures. The ATP molecule (shown in cyan/yellow sticks) and magnesium is well-defined in both chains (A&B). The binding mode of ATP and magnesium from chain B is discussed here. The adenine and ribose moiety of ATP formed key hydrogen-bonding interactions (shown in black dotted lines) with amino acid ligands from N& C lobes of HRI_KD-ΔKI_ mainly from hinge region/P-loop (highlighted in grey/yellow cartoons) are shown here. The phosphate of ATP is also coordinated by magnesium (pink sphere), water and picks up hydrogen-bonding interactions with amino acid ligands from N& C lobes

**Figure 6.**
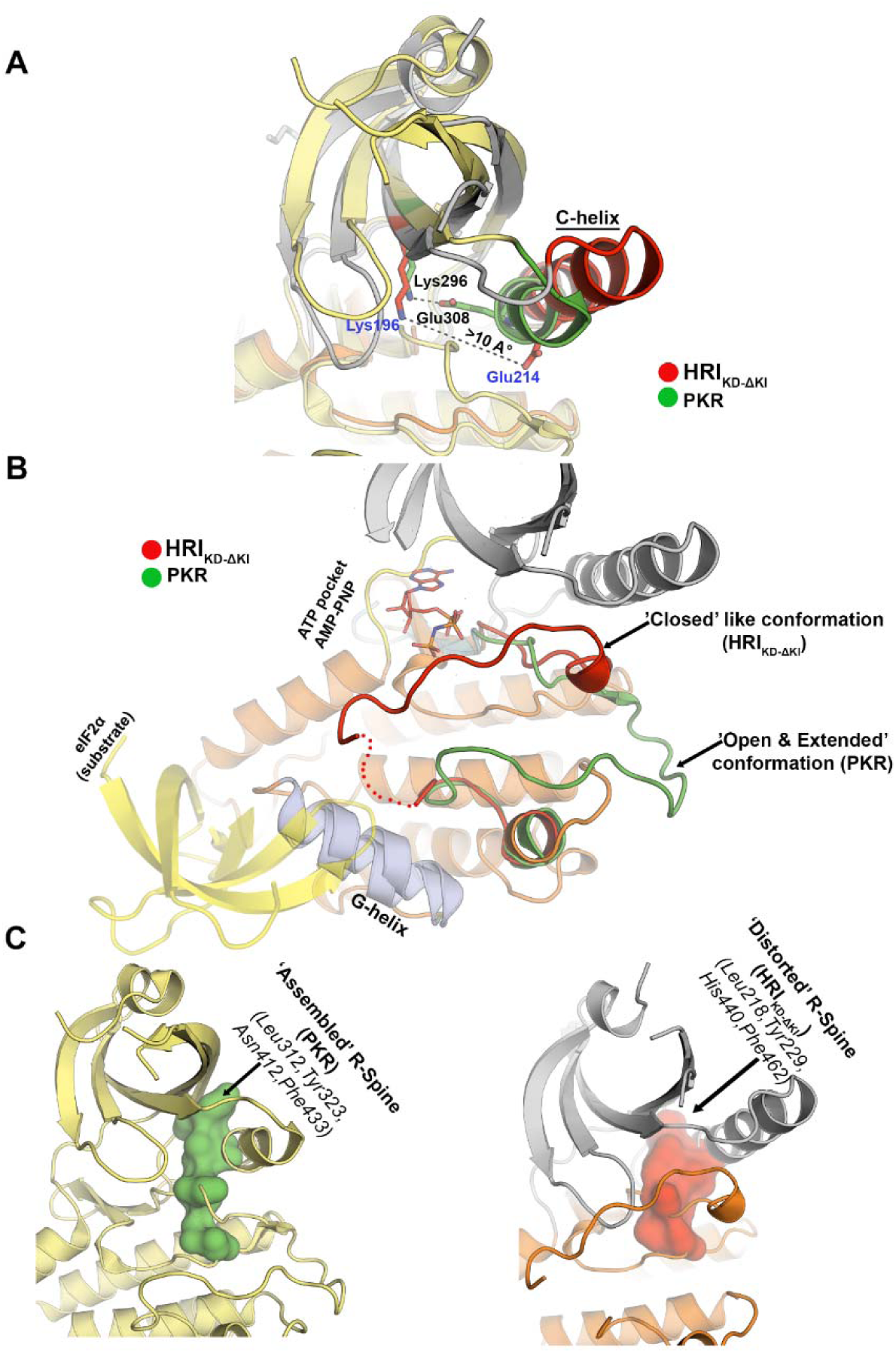
**Inactive state features observed in HRI_KD-_**_Δ**KI**_ **crystal structure.** With PKR-AMP-PNP-eIF2α complex crystal structure (PDB id:2A19) as the reference, (A) As seen generally in ‘active’ kinase structures, C-helix of PKR kinase adopts a ‘C-helix-in’ conformation with a prominent Lys296-Glu308 salt bridge (green sticks) and this is necessary for catalysis. However, in HRI_KD-ΔKI_, the C-helix adopts a ‘C-helix-out’ conformation with the outward displacement of the C-helix away from active site, which disrupts the critical Lys196–Glu214 salt bridge (shown as red sticks). (B) Unlike the active kinases where activation segment generally adopt an open and extended conformation (shows in green cartoon) e.g: for PKR resulting in productive ATP binding and substrate interaction, disordered activation segment for HRI_KD-ΔKI_ (shows in red cartoon) turns towards ATP site and thereby could hinder the ATP binding and interfering efficient binding of substrate e.g eIF2α shown in yellow cartoon (from PKR-AMP-PNP-eIF2α complex). (C) The regulatory spine (R-spine) is a key structural motif consisting of four residues named RS1-RS4, with two residues each coming from N and C-lobes. As observed in ‘active’ kinases (PKR in this case) all four residues are lined up together forming hydrophobic patch (green blob) for a catalytically functional R-spine. However, in HRI_KD-ΔKI_, the R-spine are disassembled or distorted (shown in red blob) with C-helix outward displacement resulting in an inactive state feature observed in both HRI_KD-ΔKI_ crystal structures

Close examination of the 2F_o_-F_c_ map at the ATP pocket clearly reveals strong electron density for magnesium. A single Mg²⁺ ion is observed within the structure of the HRI_KD-ΔKI_ -MgATP structure (Figure 5) in contrast to two Mg²⁺ ions typically reported in canonical PKA ternary complexes. The magnesium atom is further coordinated by oxygen atoms from the phosphates of ATP, the side chain of Asp461 and Asn447 forming a hexacoordinate-like geometry. Our observations further suggest that this Mg²⁺ ion is broadly comparable to the “Mg2” site described in the PKA ternary complexes.^53,54^

### Structural comparisons of HRI_KD-ΔKI_ with active/inactive/autoinhibited states of other structurally similar kinases

The detailed assessment of structural markers of both *apo and MgATP* bound crystal structures of HRI_KD-ΔKI_ revealed a few interesting features, which were reported earlier in inactive/autoinhibited forms of kinases including c-Src and c-AbI. The first one is that the C-helix in the N-lobe swings outwards with a conformation transition from its ‘in’ to ‘out’ state (seen in inactive c-Src, c-AbI, Figure S10). In comparison with the ‘active-state’ kinase, PKR, there is an outward rotation of 14 A° along the helix axis observed for HRI_KD-ΔKI_. This movement of the helix results in the disruption of a key salt-bridge between Lys196 and Glu214 of HRI with these residues approximately 10 AC apart (Figure 6A). This non-productive orientation prevents proper positioning of both ATP and Mg^2+^ ion needed for catalytic activity.

Secondly, in typical active-state kinase structures for PKR and PKA, the activation segment adopts an open or extended conformation (Figure 6B) that enables the productive binding of ATP and substrate simultaneously as part of the catalytic process. However, as seen in the inactive state kinases such as c-Src, Abl (Figure S10), the activation segment (spanning residues 461-500) for both HRI_KD-ΔKI_ structures, adopts an inactive, ‘closed’ like conformation that extends towards the ATP/substrate-binding site (Figure 6B and S10). The side chain of Lys196 (part of Lys-Glu bridge) forms a salt bridge with Asp461 (part of the DFG motif) and thereby the side chain of Asp461 position is moved to adopt a ‘DFG-in’ like conformation (Figure S7) and similar DFG-flip conformation (DFG-in, C-helix-out) was observed for both ‘*apo*’ and ‘MgATP’ bound HRI_KD-ΔKI_. In addition, Glu214 (part of the Lys-Glu salt bridge) is also hydrogen bonded to a backbone carbonyl of Ala465 of the activation segment. Additionally, P-loop residues (Gly177 and Gly179) interact with Asn473 of the activation loop (Figure S7). Immediately after the DFG motif, we observe a 3_10_ helix (residues 467-469) that packs against the C-helix to stabilise the C-helix out conformation (Figure S7). In short, the activation segment interacts with multiple structural elements such as Lys-Glu salt bridge, P-loop and C-helix and thereby potentially hindering access to ATP and potential substrates to the kinase domain of HRI (Figure S7 and S10).

Thirdly, the regulatory spine (R-spine), a four-residue hydrophobic core motif, is assembled and properly aligned in active kinases such as PKR (Figure 6C) and PKA, serving as a key structural hallmark required for the catalytic process. However, in HRI_KD-ΔKI_, the R-spine (His440, F462, Leu218 and Tyr229) is disrupted (Figure 6C) due to rearrangements of the C-helix and the activation segment.

## DISCUSSION

Our investigations started with exploring the stability, folding and binding kinetics of HRI_FL_. However, detailed biochemical, biophysical and structural studies were challenging, at the onset of this work, due to inconsistencies in the production of pure full-length protein, stability and folding issues. To surmount these obstacles, we designed a minimal kinase domain of HRI_FL_, named HRI_KD-ΔKI_. Deletion of the NTD domain and a 140 aa unstructured KI domain afforded a stable minimal kinase domain for HRI, that exhibited a good expression profile, producing enough pure protein in milligram quantity for biochemical, biophysical and structural studies.

Bacterially expressed HRI_KD-ΔKI_ purifies as monomer in solution in this study. The oligomeric states reported previously for HRI_FL_ varies in the range from dimer to hexamers.^46,55–58^ The recent reports ^26,46,59^ on HRI_FL_ preparations (mostly bacterially expressed) indicate HRI_FL_ exists as stable dimer in solution. The HRI_KD-ΔKI_ construct used in this study, lacking NTD, KI and CC domains, exists as a monomer in solution with an apparent MW of ∼36 kDa by gel filtration chromatography. Previous reports^46,59^ on HRI_KD_ constructs lacking an NTD and/or CC domains also reported a monomeric form with a MW of 50-70 kDa by sedimentation velocity/equilibrium analysis and demonstrated the CC and NTD subdomains are crucial and could play a potential role in HRI oligomerisation process in general. Taken together, we believe the monomeric form observed with our HRI_KD-ΔKI_ protein preparations could be due to the lack of NTD and CC domains. However, our construct lacks the KI domain also along with CC and NTD in comparison with the previously reported HRI_KD_ constructs. To our knowledge, the importance of the KI domain of HRI in oligomerisation has not yet been thoroughly characterized and its precise role warrants further investigation.

Mass spectrometric studies provided us with an overview of HRI_KD-ΔKI_ post-translational modifications characteristics and confirmed that the protein is capable of autophosphorylation. Interestingly, we observed a phosphorylated pSer409 residue (part of αD-αE loop) by mass spectrometry (Table S1) and in the HRI_KD-ΔKI_ crystal structure. Though the functional role of this residue is not studied in detail, our sequence-structure based multiple sequence alignment revealed that this Ser409 is present only in HRI compared to other related eIF2a kinase homologues (GCN2, PKR and PERK) (Figure S11). Moreover, Ser409 resides within the αD–αE loop, and this loop is longer in HRI compared to the corresponding loop regions in related eIF2α kinase homologues (Figure S11). In addition, this residue is in proximity to HRM1 heme motif (Figure 4B) which is again unique for HRI. Altogether, this warrants further investigation by mutagenesis/functional studies (biophysical/biochemical), which may help assess whether this auto phosphorylated Ser409 aids in stability, enzyme assembly and activity.

One of the main objectives of this study was to highlight the potential of HRI_KD-ΔKI_ as a viable target for structure-based drug discovery (SBDD). FTS studies, in combination with SRCD, demonstrated that the protein is stable and folded with intact secondary/tertiary features. A positive thermal stabilisation shift (ΔT of >1.0CC) from the FTS assay showed that the ATP pocket is accessible for ligand engagement. Our ligand binding assessment with reference small-molecule ligands, by FTS and SRCD assays, showed dabrafenib and GCN2iB as two promising tool compounds for HRI_KD-ΔKI_ in line with previous findings for HRI_FL_.^26^ In addition, SRCD titrations quantified the binding affinity for dabrafenib and GCN2iB with affinities ranging between 7-12 µM. Although BTdCPU is considered a bonafide activator of HRI, our structural biology efforts did not show any detectable binding apart from far-UV SRCD studies, which revealed a mild alteration in the secondary structure of HRI_KD-ΔKI_ (Figure S5C and S6). In addition, FTS studies showed a moderate increase in thermodynamic stability of the HRI upon BTdCPU addition (Figure S4, Table 1). A recent report^60^ on the mechanism of BTdCPU-dependent HRI activation postulates that it acts on HRI through a mechanism involving mitochondrial uncoupling and OMA1-DELE1 (overlapping with m-AAA protease 1- DAP3-binding cell death enhancer 1) signalling rather than direct binding with HRI. This underpins again the necessity for further assessment to ascertain whether regulation of HRI by BTdCPU occurs via transient direct binding or an indirect allosteric pathway.

Interestingly, our biophysical studies, coupled with structural biology investigations of HRI_KD-ΔKI_, provided preliminary insights into the reported key heme-binding sites (S, R, HRM1 and HRM2) of the HRI.

Thermal denaturation studies (FTS and SRCD based) showed negligible binding of hemin (HRI inhibitor ligand) to HRI_KD-ΔKI_ in line with reported thermal denaturation studies, by an FTS assay, on heme with HRI_FL_.^26,46^ Moreover, our co-crystallisation/soaking structural biology attempts to demonstrate HRI_KD-KI_-heme binding were unsuccessful with no heme bound in any of the four putative binding sites. However, in our SRCD spectral study, moderate spectral changes were detected for HRI_KD-ΔKI_ after hemin addition, at near-UV compared to far-UV, suggesting the possibility of heme-induced local tertiary structure perturbation with potential structural rearrangements in the protein (Figure S5E).

The ‘S’ heme site is not present in HRI_KD-ΔKI_ as its NTD domain is deleted in this construct. Therefore, the relevance of potential heme axial ligands such as His119/His120 lining the ‘S’ site from a three-dimensional point of view is not reported in this study. Following the ‘S’ site, HRM1 and HRM2,^3^ with a Cys-Pro (CP) motif, represent the two potential heme regulatory motifs present in HRI. Similar heme ‘CP’ motifs have been reported for heme oxygenase 2^61,62^, Bach1^63–65^, ALAS1^66,67^, JAK2 kinase^68^ and eIF2alpha kinase PK4 from Plasmodium falciparum.^69^ We mapped the HRM ligands (residues 410-415 & 552-557, Fig 4B) in our crystal structures, and they are well-defined. The multiple sequence-structure based alignment between eIF2α kinase family members (Figure S11) reveals that the HRM motif is present only in HRI although the exact role of these motifs in function regulation is not yet known.

The remaining heme site, termed the ‘R’ site, is visible in our structure, and spans the kinase domain of HRI, and comprises Cys411, His377 and His38, however, their role in heme binding events at this stage remains unclear.^34,58,70^ For example, we mapped these residues lining the ‘R’ site in our HRI_KD-ΔKI_ crystal structure, and they display prominent, well-resolved electron density for the above residues that are distributed in the N-lobe (in β5 strand) and C-lobe kinase domain of HRI. However, the spatial distribution of these ‘R’ site residues in our crystal structure showed that they are positioned far apart in the tertiary structure. This raises the question whether these residues can form a unified heme-binding pocket or contributions from additional amino acid ligands from the N and C lobe, and KI domain are still required for the successful coordination of a heme molecule. This could be one possible explanation for the absence of prominent heme binding in HRI_KD-ΔKI_, as the heme-binding pocket, ‘R’ site may not be fully formed or properly structured in HRI_KD-ΔKI_. Taken together, detailed insight at the structural level is required for a better understanding and to distinguish whether the R-site heme-sensing pocket is an intra-domain KI feature or an interlobar boundary pocket organized by the intersection of the terminal N-lobe, the hinge region, and the adjacent KI insert. The future determination of the three-dimensional structure of HRI_FL_ in complex with heme could provide detailed insight into the structural framework of heme sites of HRI.

The crystal structures of the kinase domains (KDs) of GCN2, PKR, and PERK—the remaining members of the eIF2α kinase family—have revealed well-defined dimeric configurations. In the case of PKR and PERK, a back-to-back dimeric interface mediated by key residues within the N-lobes has been observed^31,48^, whereas parallel and antiparallel dimer assemblies were reported for human and yeast GCN2, respectively.^30,47^ However, the homodimer observed in our crystal structure of HRI_KD-ΔKI_ may represent a crystallization artifact and symmetry packing. This interpretation is further supported by analytical gel filtration studies which reported that HRI_KD-ΔKI_ exists predominantly as a monomer in solution. Recent HRI dimerisation reports^46,59^ describing the detailed functional characterization of HRI_KD_ & HRI_FL_ postulated coiled-coil (CC) and N-terminal domains (NTD) as critical structural elements required in general for HRI oligomerisation process. Notably, both domains are absent in our construct, which may explain the predominantly monomeric state observed in solution. Moreover, the analysis of the crystal structure with PDBePISA supports a monomeric state with Complexation Significance Score (CSS) of 0.0.

The detailed structural comparison studies of HRI_KD-ΔKI_ revealed that, despite capturing the kinase domain of HRI in an *apo* and ATP-bound state, the protein adopts a classic “Src-like inactive” architecture, characterized by a distorted Lys-Glu salt bridge, ‘DFG-in’ motif, paired with a C-helix out conformation and a highly disordered activation loop. Interestingly, N-lobe domain movement calculations between the *apo* and ATP-bound states indicate a negligible value of 1.3C angle of N-lobe changes relative to the C-lobe upon nucleotide binding. Though the active site cleft can sterically accommodate ATP without a large rigid-body domain shift with a properly oriented DFG motif, the outward displacement of the C-helix prevents the assembly of the R-spine and disrupts the critical catalytic Lys-Glu salt bridge. Consequently, ATP binding is insufficient, thermodynamically, to transition the kinase domain beyond this trapped, pre-activation intermediate state.

This rigid structural conformation observed from our study explicitly highlights the point that our HRI_KD-ΔKI_ construct lacks a few critical conformational triggers required for full activation. Prior crystallographic studies^47^ of wild type yeast GCN2 (HRI’s closest structural homologue) reported a similar autoinhibited feature and showed activating mutations within the kinase domain, coupled with the autophosphorylation of the activation loop, serve as the indispensable conformational switches required to remodel the active site cleft for eiF2α substrate phosphorylation. As seen in GCN2, the structural features observed in our *apo* and ATP bound forms, implies that HRI might also rely on an interconnected network of regulatory input, namely, allosteric engagement of its regulatory domain especially the NTD of HRI and downstream activation loop phosphorylation to cooperatively drive and establish a catalytically competent conformation optimized for productive ATP binding followed by substrate phosphorylation. Previous biochemical evidence^71^ points to interdomain crosstalk between the NTD and the kinase domain of HRI as part of the catalytic process. Further work focusing on elucidating this intimate interdomain communication through determination of NTD-KD chimeric structures in the presence/absence of heme derivatives by X-ray crystallography and/or Cryo-EM coupled with site-directed interface mutagenesis of key heme axial ligands might help us to ascertain whether NTD has a precise role in driving the global conformational changes necessary for the transition from pre-activation intermediate to organize a productive, fully active state suited for ATP catalysis.

In conclusion, given the challenges associated with the study of HRI in its full-length form, we have successfully developed HRI_KD-ΔKI_ (minimal kinase domain of HRI). This construct has enabled the visualization of the first high resolution architecture of the core HRI kinase domain since the discovery of HRI 75 years ago, thereby solving the missing piece of the eIF2α jigsaw. This study also consolidated several unique HRI structural features, including HRMs, a non-canonical G-helix and a phosphorylation site (pSer409), which is part of a long αD-αE loop. Combined with structural findings detailing the expanded architecture of the ATP pocket framework, complementary biochemical/biophysical assays confirm the binding of key tool compounds and the presence of at least two distinct, non-competitive binding sites (GCN2iB, dabrafenib). In short, these findings provide an improved mechanistic understanding of the function of HRI, at the molecular level, and provide a firm structural foundation for future drug discovery campaigns, particularly targeting hematological malignancies.

## Materials availability

Plasmids and reagents generated in this study are available upon request to the lead contact. The sharing of plasmids/reagents may require MTA agreements.

## Data and code availability

- Atomic coordinates and structure factors for HRI_KD-ΔKI_ and HRI_KD-ΔKI_ -ATP structures have been deposited in the Protein Data Bank with accession codes, 9RHH (*apo*) (the –ATP one will be publicly available soon, under review).

## Supporting information

ESI

## ACKNOWLEDGMENTS

We thank Blood Cancer UK (grant ref# 23015) for funding this work (M.B.R., D.F.C., J.S.). We acknowledge Diamond Light Source Beamline B23 and Beamline I24 and their staff for SRCD and X-ray diffraction data collection. We thank Dr Kate Heesom and Dr Phil Lewis (Proteomics Facility, University of Bristol, UK) for phosphor-mapping studies by mass spectrometry, and Dr Joanne McCorriston and Dr Daniel Müller (Reaction Biology, Europe GmbH) for useful discussions. The BBSRC Mass Spectrometry and Proteomics Facility, University of St Andrews, is acknowledged for mass spectrometry studies. We thank BBSRC (BB/W02019X/1, E.M. P.I.) for funding a Sussex Crystallization Platform for Bioscience Discovery and EPSRC, for supporting the purchase of a single-crystal X-ray diffractometer through a Strategic Equipment Grant (EP/X013332/1) and the Wolfson Foundation for a gift towards NMR upgrades at Sussex.

## AUTHOR CONTRIBUTIONS

Conceptualization, funding acquisition, data analysis, protein preparation, characterization, crystal growth and X-ray determination, manuscript writing. M.B.R.; assay design, assay determinations and data analysis, manuscript writing, J.B.; synthesis of tool compounds, medicinal chemistry insight, data analysis, D.F.C.; X-ray determination, data analysis, S.M.R.; baculovirus protein expression, L.Z.; SRCD experiments, data analysis, manuscript writing, R.H., T.M.G., and G.S.; tool compound preparation, mass spectrometry, experimental and data analysis, R. G. M., M.S., H. H.; protein preparation insight and Xray data oversight A.W.O.; biological assay design and structural data analysis, E.J.M.; conceptualization, oversight, funding acquisition, manuscript writing, J.S.

## DECLARATION OF INTERESTS

The authors declare no competing interests.

## METHOD DETAILS

### Constructs, Expression and purification of HRI_KD-ΔKI_

Human full-length HRI (HRI_FL_) consists of amino acids ranging from 1 to 630 aa (UniprotKB ID: Q9BQI3). The truncated minimal kinase domain of HRI (HRI_KD-ΔKI_) used in this study contains selected aa ranging from 155 to 244, 374-585 with the deleted residue segment of the ‘KI’ domain extending between 245-373 amino acids. The entire ‘KI’ is replaced with a two-residue flexible linker ‘GS’ for this study. The codon optimised bacterial expression vectors coding for N-term GST truncated HRI with minimal kinase domain (HRI_KD-ΔKI_) protein used in this study were purchased from Genscript Biotech (UK) Limited.

### Expression and purification of HRI_KD-ΔKI_

pGEX-GST-3C- HRI_KD-ΔKI_ plasmid was transformed with BL21(DE3) (New England Biolabs) as host cells and plated on Luria Broth (LB) -agar containing an appropriate antibiotic (100 μg/mL Ampicillin) and incubated overnight at 37 CC. A ’10 mL’ bacterial starter culture was used to inoculate 1 L Turbo-broth (TB) (Cat no: MD12-104; Molecular Dimensions) in a 2 litre Erlenmeyer flask supplemented with 100 µg/mL Ampicillin. The cultures were then allowed to grow at 37 CC under agitation at 200 rpm until O.D. reached 1.5-2.0 for TB. HRI_KD-ΔKI_ protein expression was induced with the addition of 0.5 mM isopropyl-D-thiogalactopyranoside (IPTG) followed by further incubation of the culture in the shaker overnight at 18 CC for 15-16 h.

The pellets from 1 L of cells were resuspended using 2 mL ‘chilled’ Buffer A (50 mM HEPES pH 7.5, 750 mM NaCl, 0.5 mM tris(2-carboxyethyl) phosphine hydrochloride (TCEP), 5% Glycerol) per gram of cell pellet. The lysate was further supplemented with 150 µL Turbo DNAase (Cat no: AM2239; Thermofisher Scientific) and six EDTA-free protease inhibitor cocktail tablets from Roche Diagnostics (Cat no: 11836170001). The cells were disrupted by sonicating cells using sonicator (Fisherbrand Sonic Dismemebrator FB-505). Debris and insoluble materials were removed by centrifugation in a Beckman Coulter, rotor ID 25.5 (35,000 xg, 4 CC for 1 h). The supernatant, after centrifugation, filtered through a 0.45 µM sterile syringe filter (Fisher Scientific) before treatment with affinity resins.

A two-step purification protocol was employed for the purification of HRI_KD-ΔKI_ using GE AKTA Purifier 10 FPLC system. The first one was a GST-affinity purification with incubation of above processed supernatant with Glutathione Sepharose 4B resin (Cat. no: GE17-0756-01; Cytiva) preequilibrated with Buffer A for 2 h with rotation at 4 CC with a Econo-Chromatography gravity column (Bio-Rad) for the initial capture of GST-fusion HRI_KD-ΔKI_. The unbound proteins were then removed by washing resin with 100 mL Buffer A. Following the washing step, the glutathione resin slurry with bound GST-fusion HRI_KD-ΔKI_ was mixed with in-house HRV-3C protease (150 µL of ∼2mg/ml) for GST tag cleavage removal reaction in a 15- or 50 mL falcon tube at 4 CC, rolling overnight. The following day, HRI_KD-ΔKI_ protein fractions were eluted with Buffer A. Fractions containing HRI_KD-ΔKI_ were identified by SDS-PAGE followed by pooling of selected ‘HRI_KD-ΔKI_’ fractions and concentrated using centrifugal ultra-filtration with a 10 kDa MWCO Vivaspin concentrator (Cat no: VS2002; Sartorius) according to the requirements for the second step of purification. The second step, as well as the final polishing purification step for HRI_KD-ΔKI_, involved loading HRI_KD-ΔKI_ protein fractions to a HiLoad 16/60 Superdex 200 size exclusion column (Cytiva), pre-equilibrated with Buffer B (25 mM HEPES pH 7.5, 250 mM NaCl, 0.5 mM TCEP, 5% glycerol). The fractions from this purification step were identified by SDS-PAGE and concentrated to a final desired concentration of ∼15 mg/mL to produce pure soluble HRI_KD-ΔKI_ on a large scale (up to 15 mg per batch) and stored at –80 °C until required. The identity of the final purified proteins generated in this study was confirmed by performing trypsin digests on Commassie stained gel bands followed by mass spectrometric analysis in each case (BSRC Mass Spectrometry and proteomics Facility, University of St. Andrews, UK). The overall molecular mass of HRI_KD-ΔKI_ was further assessed with in-house LC-MS analysis using an Equan 850 HPLC coupled to an Orbitrap Exploris 480 (Thermo Fisher Scientific) equipped with a HESI ion source. Briefly, HRI_KD-ΔKI_ protein was buffer exchanged to 100 mM ammonium acetate buffer using Sartorius Vivaspin 500 centrifugal concentrator according to the manufacturer’s instructions. The final concentration of the HRI_KD-ΔKI_ after buffer exchange was 5 µM for the MS studies. Data was analysed using BioPharma Finder 5.2 (Thermo Fisher Scientific).

### Phosphorylation status of HRI_KD-ΔKI_ by mass spectrometry

The distinct gel bands related to HRI_KD-ΔKI_ after SDS-PAGE were carefully excised from the gel and sent to the University of Bristol Proteomics facility for HRI_KD-ΔKI_ phospho-sites identification. The gels were subjected to in-gel tryptic digestion using a DigestPro automated digestion unit (Intavis Ltd). The resulting peptides were fractionated using an Ultimate 3000 nano-LC system in line with an Orbitrap Fusion Lumos mass spectrometer (Thermo Scientific). All spectra were acquired using an Orbitrap Fusion Lumos mass spectrometer controlled by Xcalibur 3.0 software (Thermo Scientific) and operated in the data-dependent acquisition mode. The raw data files were processed and quantified using Proteome Discoverer software v2.1 (Thermo Scientific) and searched against the UniProtKB *Escherichia coli* (strain BL21-DE3) database (downloaded June 2024; 4156 sequences) and a database containing the HRI_KD-ΔKI_ sequence using the SEQUEST HT algorithm. Peptide precursor mass tolerance was set at 10 ppm, and MS/MS tolerance was set at 0.6 Da. Search criteria included oxidation of methionine (+15.995 Da), phosphorylation of serine, threonine and tyrosine (+79.966 Da), acetylation of the protein N-terminus (+42.011 Da) and methionine loss plus acetylation of the protein N-terminus (−89.03 Da) as variable modifications and carbamidomethylation of cysteine (+57.021 Da) as a fixed modification. Searches were performed with full tryptic digestion and a maximum of 2 missed cleavages were allowed. The reverse database search option was enabled, and all data were filtered to satisfy false discovery rate (FDR) of 5%.

### Kinase assays for HRI_KD-ΔKI_ and HRI_FL-RB_

A radiometric, filter plate-based assay was used by Reaction Biology to directly measure HRI protein kinase activity. The full-length HRI (Cat no: RBE product #0444-0000-1) used in this assay is provided by Reaction Biology and hereafter referred as ‘HRI_FL-RB_’. Reactions were performed at 30 °C for 60 min using 40 nM HRI (HRI_FL-RB_ or HRI_KD-ΔKI_), and a range of ATP concentrations up to 30 µM (using a mixture of [γ-³³P]ATP, (Hartmann-Analytic, Cat# FF301T), unlabelled ATP (Promega, Cat no: V915B-C) and substrate casein (Sigma-Merck Cat no: C4765) at 2µg/50µL. Reactions were assembled in 96-well polypropylene plates (ThermoFisher, Cat no: 442587) in a total volume of 50 µL assay buffer containing: 70 mM HEPES-NaOH pH 7.5, 3 mM MgCl₂, 3 mM MnCl₂, 3 µM sodium orthovanadate, and 1.2 mM DTT. Reactions were stopped by the addition of 200 µL of 10% H₃PO₄ and transferred to pre-wetted MSFC/PH filter paper (Merck, Cat no: MSFCN6B). The filters were washed with 10% H₃PO₄, dried for 30 min at 45 °C, and scintillation cocktail (50 µL) was added. Radioactivity was measured using a microplate scintillation counter (PerkinElmer). Radioactivity retained on the filter (representing phosphorylated substrate only) was quantified as counts per minute (cpm), providing a direct measure of kinase activity. Reaction velocities were calculated as pmol phosphate incorporated per minute per µg enzyme (pmol/µg/min). Data were plotted as a function of ATP concentration and visualised in GraphPad Prism 10 by linear connection of data points.

### FTS assays for HRI_KD-ΔKI_

The fluorescence-based thermal shift assays for thermal denaturation studies for *apo* and ligands/substrates incubated forms for HRI_KD-ΔKI_ were performed in 96 well plates using a Roche LightCycler 480 II. The excitation and emission wavelengths for the detection of protein unfolding were 465 and 580 nm respectively. The standard denaturation reaction involved temperature gradient ranging between 20 to 85 CC with the heating ramp rate of 4.4 CC/s and cooling down rate of 2.2 CC/s. The data acquisition was 20 data measurements taken per degree Celsius. Purified protein (HRI_KD-ΔKI_ was diluted in Buffer B (25 mM HEPES pH 7.5, 250 mM NaCl, 0.5 mM TCEP, 5% glycerol) according to the requirements. The final volume of protein alone-dye mixture for each well was 20 µL with concentration of 1.87 µM and 5x SYPRO Orange dye whereas for protein/ligands incubated forms-dye mixture, the ligands-protein ratio was approximately 10:1 (25 µM-2.5 µM) with the DMSO final concentration of <1%. Following the sample addition to the wells, the plates were sealed using LightCycler^®^ Sealing Foils before running assays. The spectral measurements of each sample were recorded as either three or six scan replicates using Roche LightCycler 480 II. The raw data for melting temperature (T_m_) curves from Roche Light Cycler 480 were processed using a modified Boltzmann model equation described previously.^35,72^ The representative thermal denaturation profile spectra for *apo* and ligand incubated forms for HRI_KD-ΔKI_ were generated by plotting the normalised melting curve using Graphpad Prism 10 version.

### Characterisation of HRI_KD-ΔKI_ folding and ligand binding by SRCD

SRCD spectra were acquired on beamline B23 (Diamond Light Source) using nitrogen flushed ChirascanPlus and B23 endstation-A spectrometer. HRI_KD-ΔKI_ samples (∼1CmgCmL⁻¹) typically prepared in 10 mM HEPES pH 7.5, 75 mM NaCl, 0.25 mM TCEP were analysed in far-UV (190–260 nm) for secondary structure and near-UV (250–350 nm) for tertiary structure using 0.1 mm and 1 cm pathlengths cell respectively with 1nm bandwidth and 1 sec integration time at 20 °C.

Ligands (dabrafenib, GCN2iB, BTdCPU, hemin) were added at a 1:2 protein:ligand stoichiometry, unless otherwise specified. The thermal denaturation studies by SRCD were monitored by measuring far UV CD spectra between 20-90°C. Data were fitted with a Boltzmann model to determine unfolding transitions at 209Cnm over 20–90 °C. Ligand-induced stabilisation was assessed from relative shifts in signal profiles and high-temperature plateau behaviour. The near-UV SRCD HRI_KD-ΔKI_ -ligand titrations were performed by monitoring ellipticity changes (∼338–355 nm) as a function of ligand concentration. Binding curves were fitted using a single site/Hill model to determine apparent dissociation constants (K_d_). Values reported represent fitted parameters from replicate datasets, including independent repeat experiments.

### Crystallisation, data processing and structure determination of HRI_KD-ΔKI_

Purified HRI_KD-ΔKI_ (in 25 mM HEPES pH 7.5, 250 mM NaCl, 0.5 mM TCEP, 5% glycerol) was concentrated to ∼15 mg/mL using a centrifugal membrane concentrator (Vivaspin, 10 kDa MW cut off). A broad initial HRI_KD-ΔKI_ crystallisation screening was performed with a protein crystallisation robot dispenser, Oryx8 from Douglas Instruments Ltd by screening around ∼400 crystallisation conditions. The tested crystallisation conditions are from commercial 96-reagent crystallisation screens kits such as JCSG Plus (Cat no: MD1-40), PACT Premier (Cat no: MD1-36), Structure (Cat no: MD1-30) (from Molecular Dimensions Ltd), and PEG_Ion (Cat no: HR2-139 from Hampton Research). Each droplet for crystal formation was prepared by mixing 0.2 µL of HRI_KD-ΔKI_ (12 mg/mL) with 0.2 µL of reservoir well solution at room temperature with sitting-drop vapor-diffusion method and equilibrated against a 50 µL reservoir well solution. From ∼400 crystal conditions screened, a few promising leads (5 to 10 hits) started to be observed after three weeks incubation at 4 °C showing needle clusters, cubic shaped crystals etc. Crystals of ‘apo’ HRI_KD-ΔKI_ were carefully harvested from the crystal drop and mounted using CryoLoops (Hampton Research) and cryocooled in liquid nitrogen before the X-ray diffraction data collection. The cryoprotection was done with respective crystal conditions supplemented with a final 20% glycerol as cryoprotectant.

The X-ray diffraction data for the protein crystals of HRI_KD-ΔKI &_ HRI_KD-Δ_KI-MgATP were collected on macromolecular crystallography Beamline I24 & I04 respectively (Diamond Light Source, UK) at 100 K. Data processing for the determination of HRI_KD-ΔKI_ crystal structure was performed using programmes provided in the CCP4 suite^73^ and PHENIX.^74^ The crystal structure for HRI_KD-ΔKI_ was solved by molecular replacement, using the CHAINSAW & PHASER program implemented in the CCP4 package. The closest structural homologue and another prominent member of eIF2α kinase family, GCN2 from *Saccharomyces cerevisiae*^47^ (PDB code:1ZYC) was used as the search model template to look for possible packing solutions for HRI_KD-ΔKI_ using PHASER. The refinement of the HRI_KD-ΔKI_ structure solution was completed using a PHENIX and BUSTER package. The manual building of the HRI_KD-ΔKI_ model and the addition of water/ligand molecules (manually and/or automatically) to HRI_KD-ΔKI_ model during the iterative refinement steps, were accomplished using COOT^75^, and PHENIX, where appropriate. The quality of the final model was verified using Molprobity.^76^ The data collection and structure refinement statistics, generated using Table 1 module of PHENIX, are summarised in Table S2.

A PDBeFold^77^ web server and DALI^78^ server were mainly used for structure-based alignments between HRI_KD-ΔKI_ and other eIF2α kinase family members as well as for searching for similar structures against HRI_KD-ΔKI_ as the search probe. The electron density map generation (for phosphorylated residue S409 and ATP), graphic representation and interpretation of the structures for HRI_KD-ΔKI_ used in this study were performed using PYMOL (The PyMOL Molecular Graphics System, Version 3.1.6.1 Schrödinger, LLC.).

## Notes

### Competing Interest Statement

The authors have declared no competing interest.

