## Supplementary material for "Structure-function studies of HRI_KD-ΔKI_, a Minimal Kinase Domain of Human Heme-Regulated Inhibitor Kinase": ESI

**Document S1. Figures S1–S11, Tables S2**

**Table S1. List of phosphorylation sites identified for $HRI_{KD-\Delta KI}$ by mass spectrometry**

### SUPPLEMENTARY FIGURES

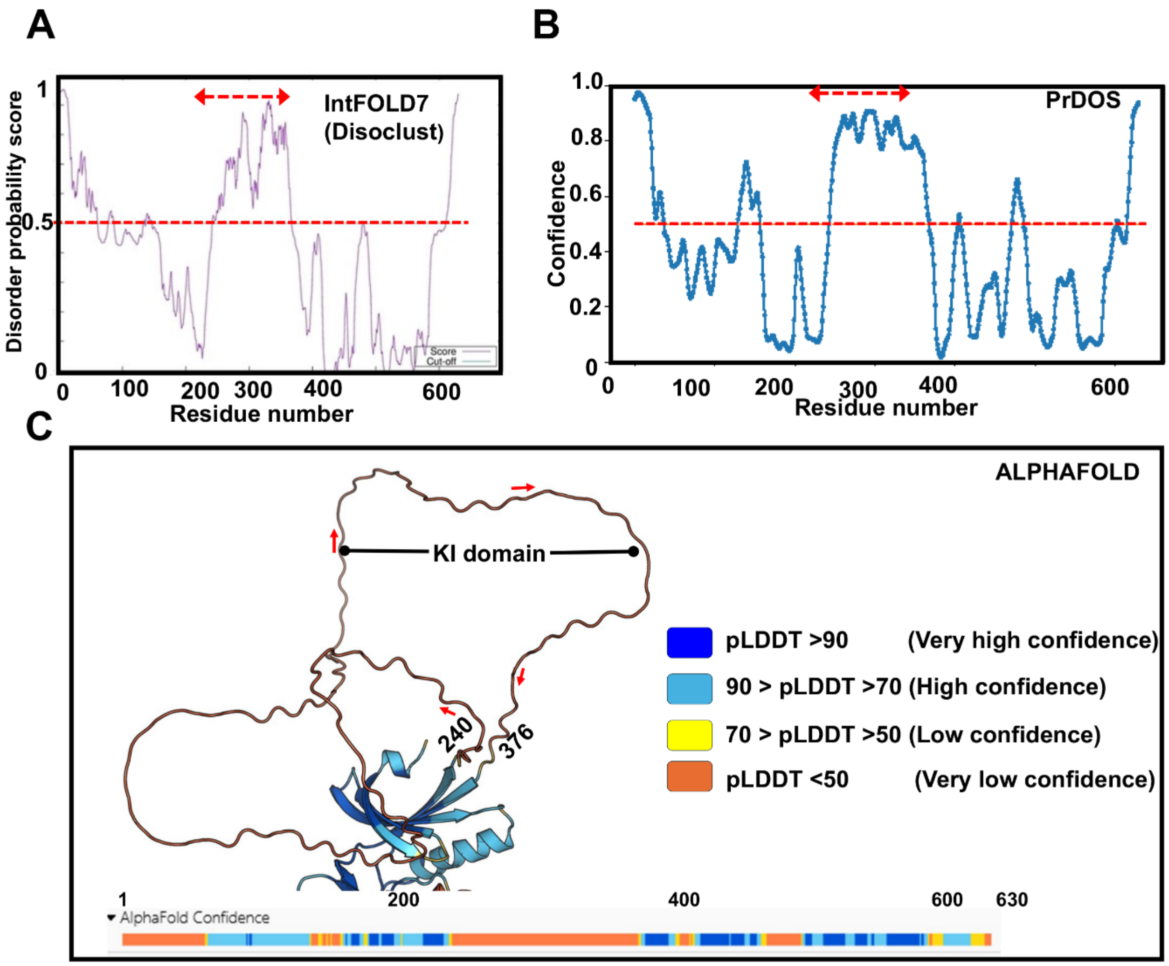

**Figure S1. In silico-based prediction of disordered regions for HRI.** A bioinformatics-based assessment of the protein sequence of HRI as part of designing optimal protein construct for HRI structure/function studies was performed using protein domain/disorders web-based tools. (A and B) Disorder probability plot of HRI from PrDOS and IntFOLD7 web-based servers, in consensus clearly suggests a disordered/unstructured region between amino acids (aa) 200 to 400 especially between 220 to 375 aa. The red dashed lines at cut off value of 0.5 refer to disorder probability/confidence threshold line and residues above these lines are predicted to be disordered. (C) Alphafold predicted structural model for HRI (Alphafold accession ID: AF-Q9BQI3-F1-v4). For clarity, the N-lobe and KI domain of HRI are shown here. predicted local distance difference test (pLDDT) from Alphafold is per-residue measure of model confidence scaled from 0 to 100. The higher the score indicates higher confidence with more accurate prediction of models. The pLDDT score <50 is typically assigned by Alphafold when there are flexible or intrinsic disordered regions in protein. For HRI, Alphafold suggests pLDDT <50 for region 240-376 postulating the KI domain is a flexible or unstructured region.

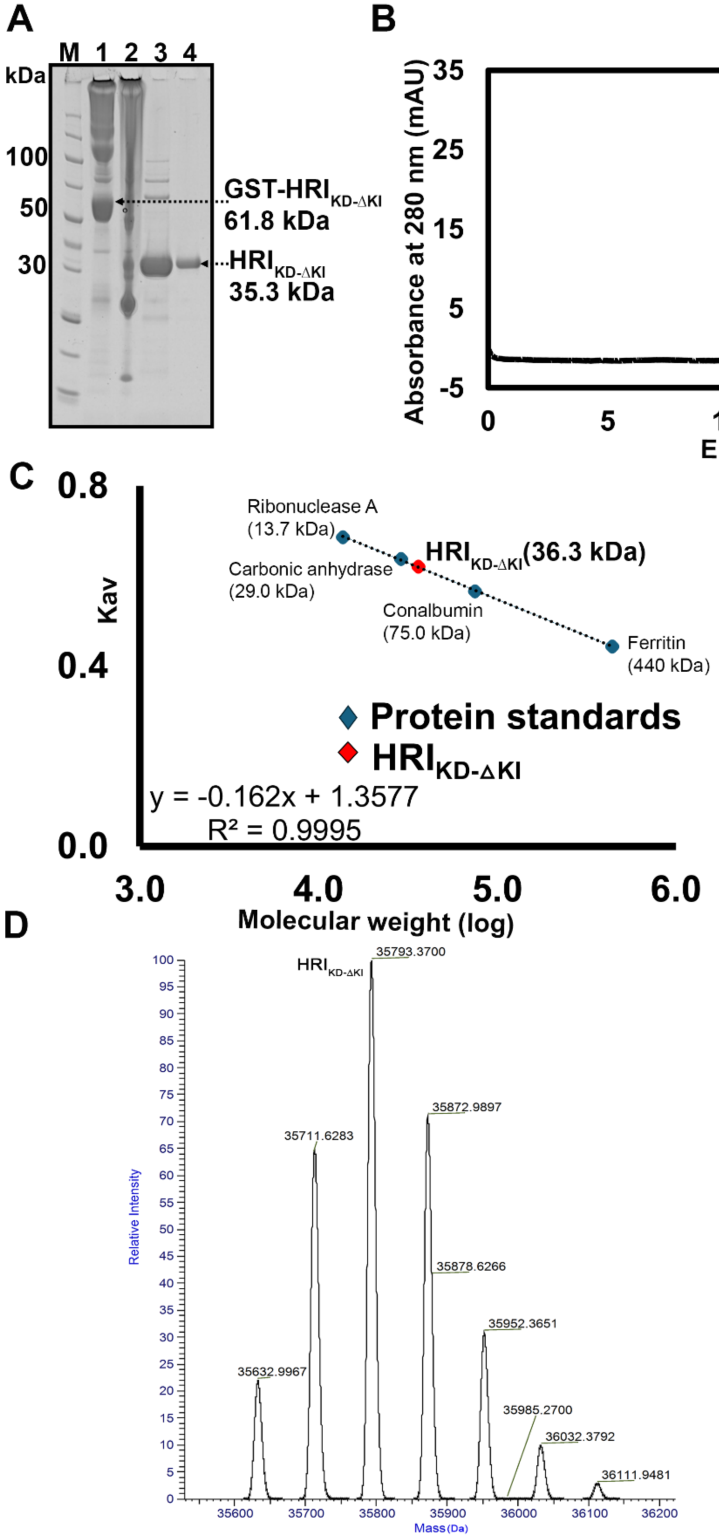

A
M 1 2 3 4
kDa
100
50
30
GST-HRI KD-ΔKI
61.8 kDa
HRI KD-ΔKI
35.3 kDa
B
Absorbance at 280 nm (mAU)
HRI KD-ΔKI
Elution (ml)
C
Kav
Ribonuclease A (13.7 kDa)
Carbonic anhydrase (29.0 kDa)
HRI KD-ΔKI (36.3 kDa)
Conalbumin (75.0 kDa)
Ferritin (440 kDa)
Protein standards
HRI KD-ΔKI
y = -0.162x + 1.3577
R² = 0.9995
Molecular weight (log)
D
HRI KD-ΔKI
35793.3700
35711.6283
35872.9897
35878.6266
35952.3651
35632.9967
35985.2700
36032.3792
36111.9481
Relative Intensity
Mass (Da)

**Figure S2. Protein production, oligomeric state calculations for $HRI_{KD\text{-}\Delta KI}$.** (A) For $HRI_{KD\text{-}\Delta KI}$ protein, a prominent band at an apparent molecular weight of 40 kDa was observed from SDS-PAGE analysis for all stages of purification matched with the expected size of 35.3 kDa. Lanes 3 and 4 correspond to $HRI_{KD\text{-}\Delta KI}$ protein after GST-affinity capture and size-exclusion chromatography respectively. (B) The oligomeric state of $HRI_{KD\text{-}\Delta KI}$ was assessed using size-exclusion chromatography by applying protein to Superose 6 10/300 column (Cytiva). The elution profile for the $HRI_{KD\text{-}\Delta KI}$ is shown here. (C) The molecular weight of $HRI_{KD\text{-}\Delta KI}$ was calculated by comparing its elution volume with a few representative standard proteins from gel filtration calibration kits LMW, HMW; Cat no: 28403841& 28403842). Gel filtration analysis revealed $HRI_{KD\text{-}\Delta KI}$ with an apparent MW of 36.3 kDa against a calculated MW of 35.3 kDa suggesting $HRI_{KD\text{-}\Delta KI}$ exists predominantly as a monomer in solution. The difference of ~1 kDa observed for $HRI_{KD\text{-}\Delta KI}$ could be accounted for the possible post-translation modifications with multiple phosphorylations. The calibration curve is plotted with a gel-phase distribution coefficient (Kav) versus logarithm of the molecular weight (log). The red and blue diamond refers to Kav for $HRI_{KD\text{-}\Delta KI}$ and protein standards used in this study. (D) Deconvoluted mass spectrum for $HRI_{KD\text{-}\Delta KI}$

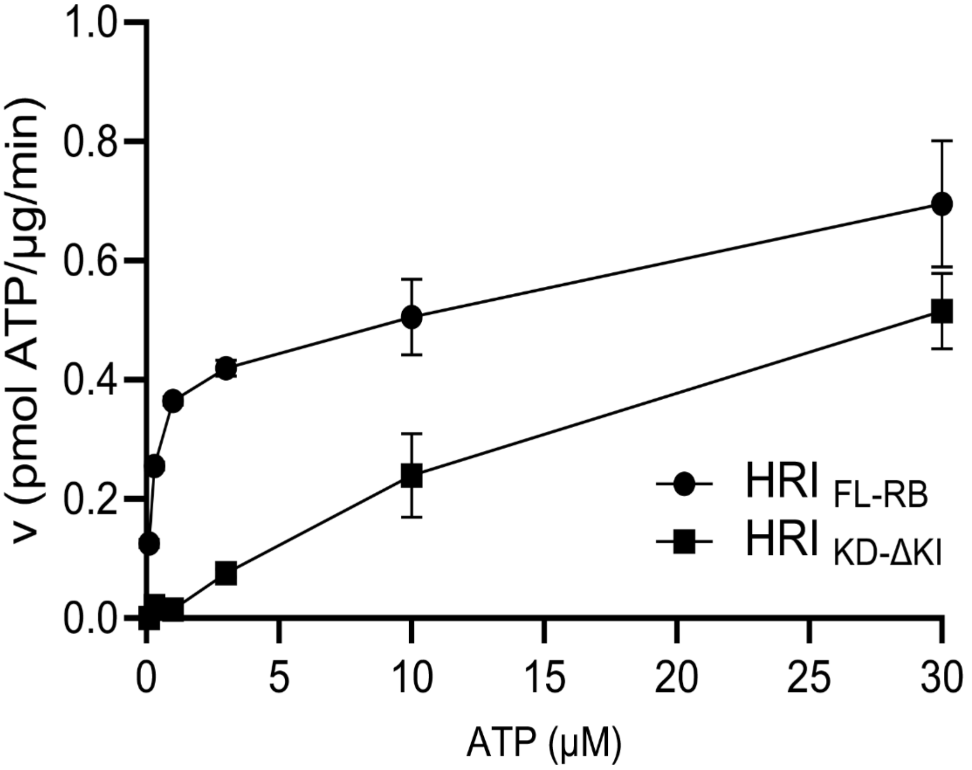

**Figure S3. HRI Enzyme Activity.** Reaction progress curves showing kinase activity of both full-length HRI ($HRI_{FL-RB}$) and $HRI_{KD-\Delta KI}$ measuring direct substrate phosphorylation using radiometric filter-based assay.

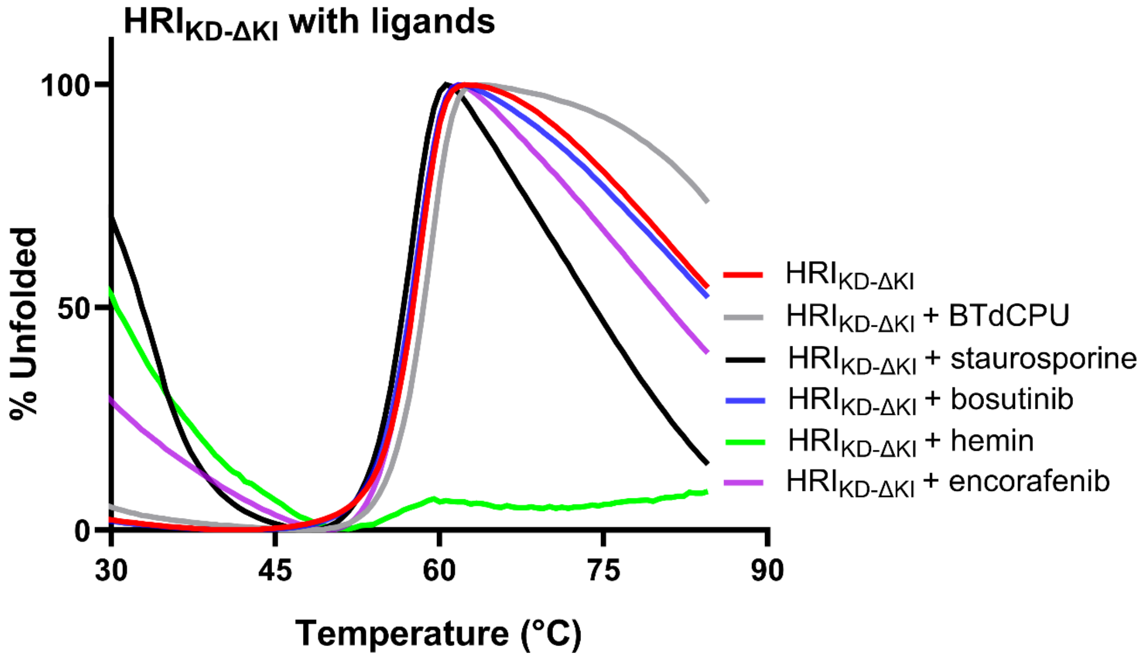

**Figure S4. FTS assay for $HRI_{KD\text{-}\Delta KI}$ – reference small-molecule ligands.** The ligand incubated forms of $HRI_{KD\text{-}\Delta KI}$ were subjected to FTS analysis as described in the methods. The displayed profile is the representative thermal denaturation profile for *apo* and ligand incubated forms of $HRI_{KD\text{-}\Delta KI}$. Apo $HRI_{KD\text{-}\Delta KI}$ exhibited a single-phase transition denaturation profile with the $T_m$ ~58.0 ºC depicted as red line. BTdCPU showed a moderate positive TM shift (grey line) of 0.8 ºC whereas remaining ligands such as staurosporine, bosutinib, encorafenib and hemin (HRI inhibitor ligand), binding was either not detected or a negative TM shift from the data analysis The spectrum shown represents the final averaged data from three consecutive scans (n= 3 scans)

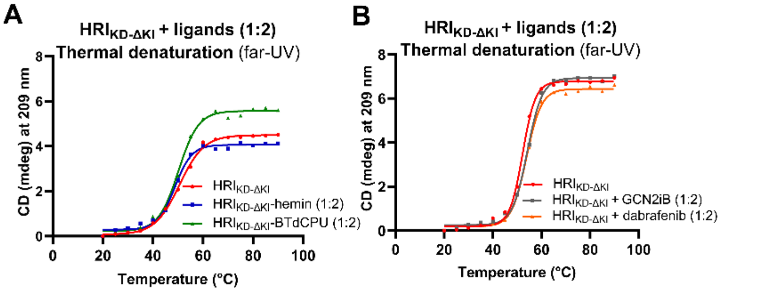

| Sample | Tm | Sample | Tm |
|---|---|---|---|
| **HRI$_{KD\text{-}\Delta KI}$** | 50.61 °C *(49-83 - 51.38 °C)* | **HRI$_{KD\text{-}\Delta KI}$** | 52.24 °C *(51.65 - 52.82 °C)* |
| **HRI$_{KD\text{-}\Delta KI}$ + hemin** | 48.74 °C *(47.63 - 49.80 °C)* | **HRI$_{KD\text{-}\Delta KI}$ + GCN2iB** | 54.17 °C *(53.59 - 54.73 °C)* |
| **HRI$_{KD\text{-}\Delta KI}$+ BTdCPU** | 50.18 °C *(49.22 - 51.11 °C)* | **HRI$_{KD\text{-}\Delta KI}$ + dabrafenib** | 53.82 °C *(53.27 - 54.36 °C)* |

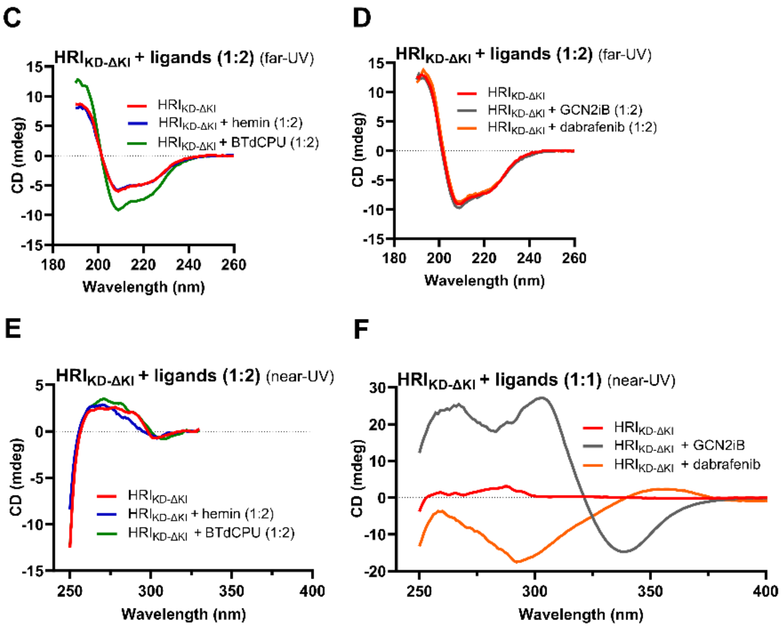

**Figure S5. SRCD thermal denaturation and Far-UV/Near-UV CD spectra studies of** HRI$_{KD\text{-}\Delta KI}$ in the absence and presence of ligands. (A-B) SRCD thermal denaturation curves of HRI$_{KD\text{-}\Delta KI}$ in the absence and presence of ligands, monitored at 209 nm. Ligand-dependent stabilisation is observed, with BTdCPU showing the strongest effect with elevated ellipticity retained across the full temperature range (60–90 °C). GCN2iB and dabrafenib were shown moderate stabilisation with ΔTm shift of +1-2 ºC. The thermal stability of HRI$_{KD\text{-}\Delta KI}$ was reduced after the addition of hemin suggesting of weak structural perturbation in the case of heme. The melting profile represents a Boltzmann sigmoidal fit of the mean CD signal monitored at 209 nm across a temperature gradient using GraphPad Prism. Each data point represents the average of four consecutive scans (n = 4) acquired at each temperature. The melting temperature (Tm) was determined by non-linear regression, and parameter uncertainty is reported as the 95% confidence interval of the fitted parameter. (C-D) Far-UV SRCD spectra (190–260 nm) of HRI$_{KD\text{-}\Delta KI}$ in the absence and presence of ligands. The observed spectral features indicate a mixed α/β fold with ligand-induced secondary structural perturbations. (E) Near-UV SRCD spectra (250–330 nm) showing ligand-induced changes in tertiary structure arising from aromatic residue environments for hemin and BTdCPU. (F) Near-UV SRCD spectra (250 - 450 nm) demonstrating Induced Circular Dichroism (ICD) effect for GCN2iB and dabrafenib with CD spectra changes observed at 338 and 355 nm respectively. The spectrum shown in all four cases (C-F) represents the final averaged data from four consecutive scans (n= 4 scans)

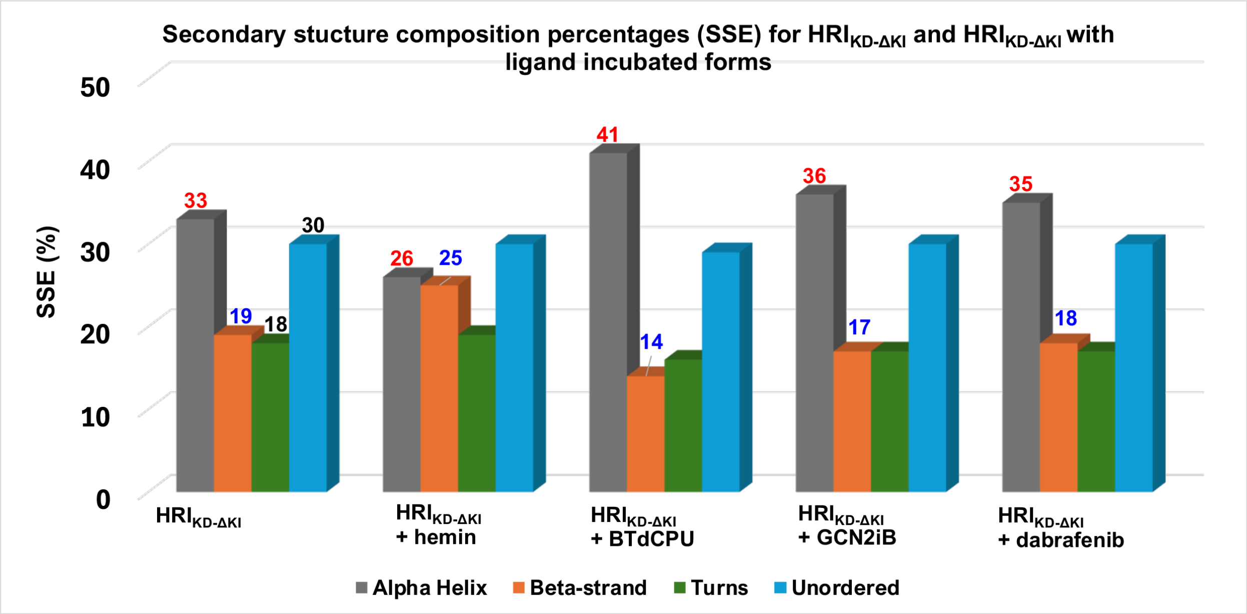

**Figure S6. Secondary structure estimation (SSE) of** $HRI_{KD\text{-}\Delta KI}$ in the absence and presence of reference small-molecule ligands. The relative percentages of secondary structural elements were calculated for all samples. The α-helical and β-sheet contents are highlighted in red and blue, respectively, for each sample.

Close-up view of activation segment of HRIKD-ΔKI

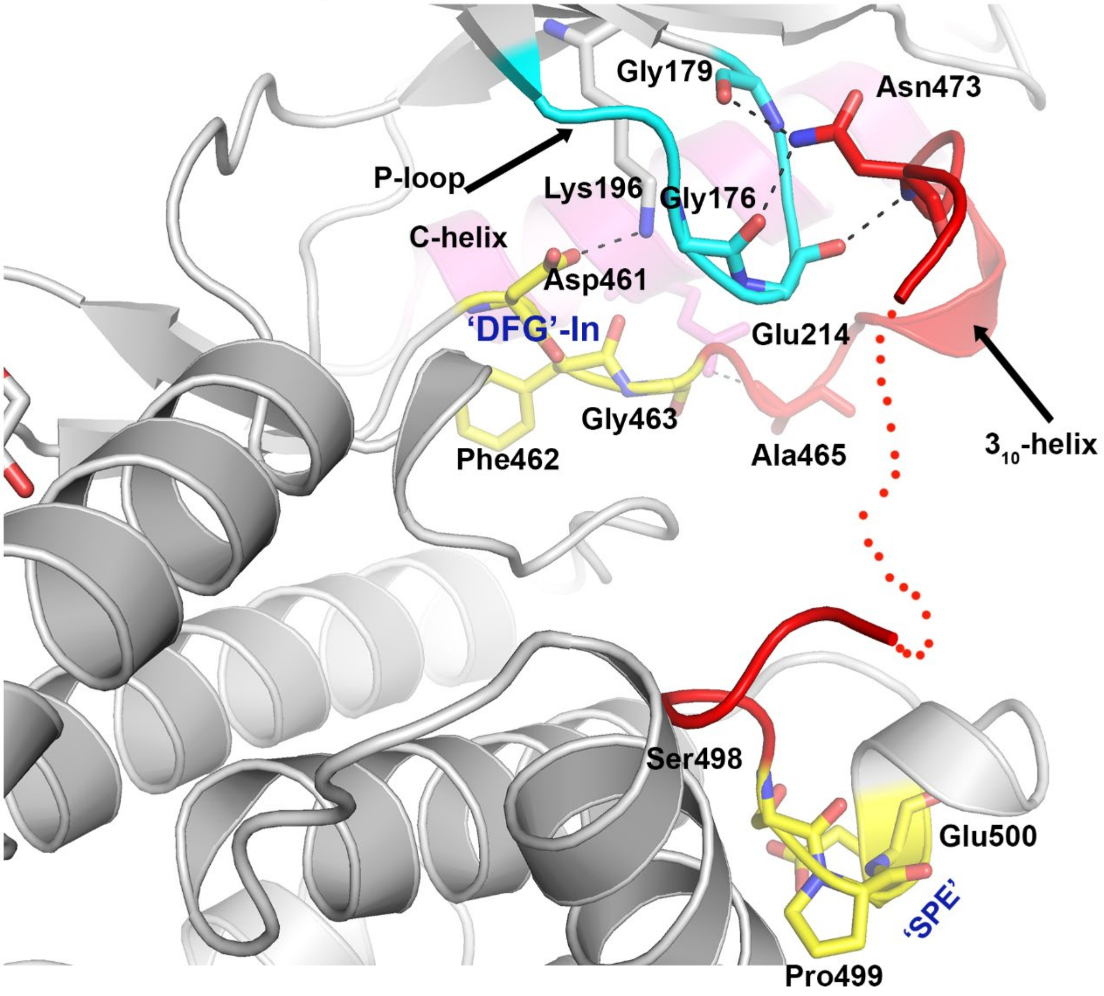

**Figure S7. Close-up view of the activation segment of $HRI_{KD\text{-}\Delta KI}$.** Both apo and MgATP bound form of $HRI_{KD\text{-}\Delta KI}$ shows a 'DFG' in conformation as generally seen in active kinases. However, the activation segment adopts a 'closed' like conformation backed by interactions with key structural elements from both N- and C-lobes. Lys196 and Glu214 (part of Lys-Glu salt bridge) is hydrogen bonded to Asp461 of 'DFG' motif and Ala465 from activation loop. Similarly, Asn473 of activation loop also picks up few interactions from amino acid ligands of P-loop. Another interesting feature observed is, presence of $3_{10}$ helix which packs against C-helix and thereby stabilising the C-helix 'Out' conformation. The activation segment extending from 461-500 is shown in red cartoon. For simplicity and consistency, $HRI_{KD\text{-}\Delta KI}$ is shown here and aa residues 479 to 492 were disordered and not modelled in this structure and it is shown as red dotted lines.

**A**

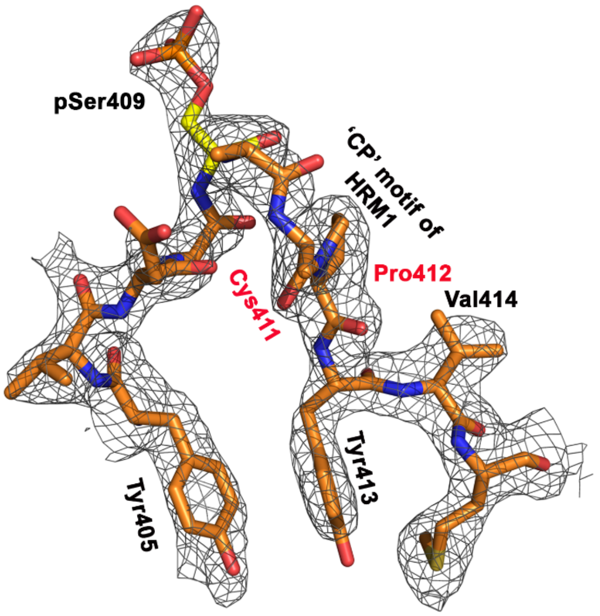

**B**

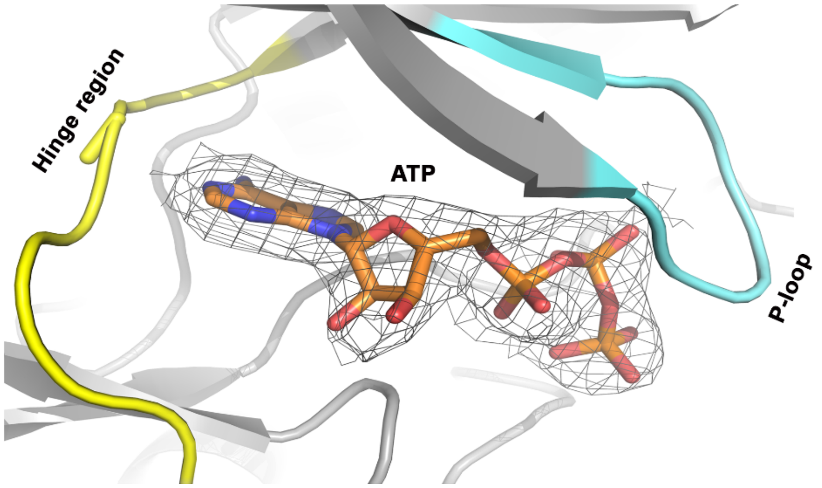

**Figure S8. Electron density maps of pSer409 and bound ATP** (A) $2F_o$-$2F_c$ electron density map contoured at σ1.0 focusing on the αD-αE linker. The region between aa 405 to 414 of $HRI_{KD-\Delta KI}$ is displayed with phosphorylation observed at Ser409. The extra unique density is clearly visible for the phosphate group. The interesting observation is, 'CP'motif which is part of HRM1 is in the vicinity of this pSer409 (B) Close-up view of the ATP-binding pocket of $HRI_{KD-\Delta KI}$- MgATP crystal structure with $2F_o$-$2F_c$ electron density map showing bound ATP contoured at σ1.0.

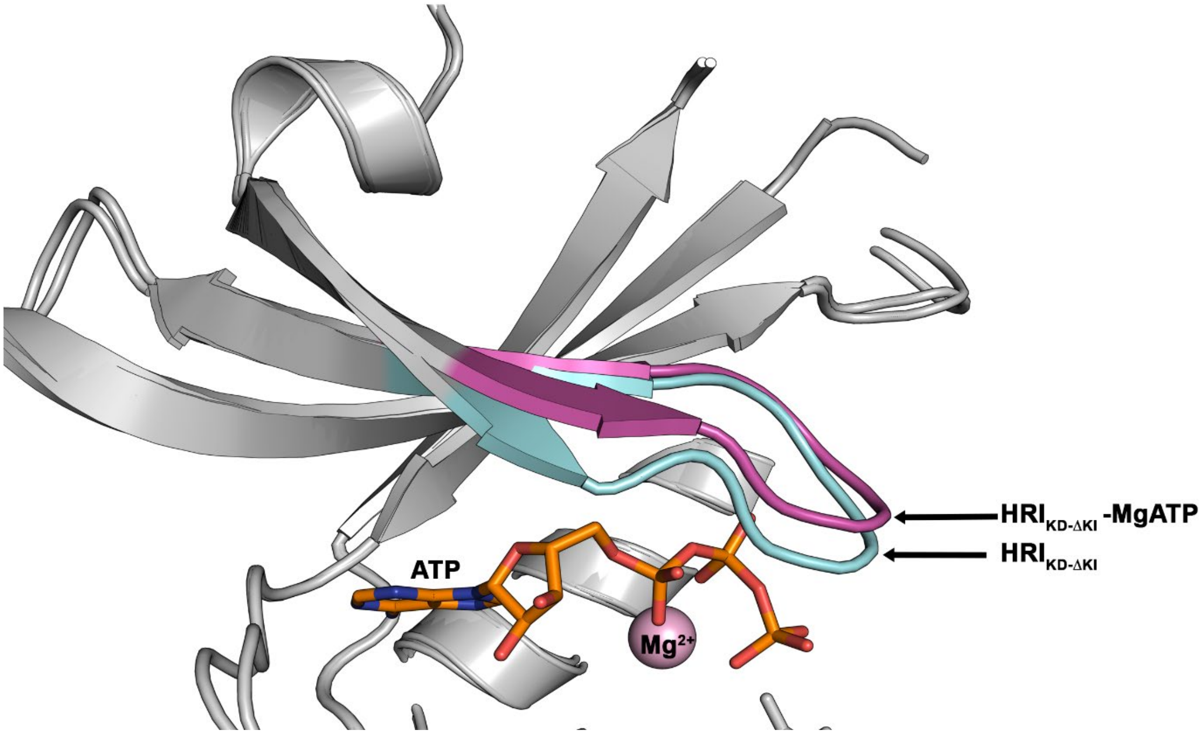

**Figure S9. Close-up view of P-loop of HRI$_{KD-\Delta KI}$.** For clarity, a close view of the N-lobes of HRI$_{KD-\Delta KI}$ (grey color) and HRI$_{KD-\Delta KI}$ -MgATP crystal structures (shown in grey color) with the P-loop of HRI$_{KD-\Delta KI}$ (cyan color) with HRI$_{KD-\Delta KI}$ -MgATP (magenta) is displayed here. As seen in the figure, P-loop of both apo and ATP bound forms of HRI$_{KD-\Delta KI}$ is well-ordered in our structure. The P-loop of HRI$_{KD-\Delta KI}$ -MgATP adopts a slightly 'open' conformation in comparison to the *apo* form of HRI$_{KD-\Delta KI}$ to accommodate ATP.

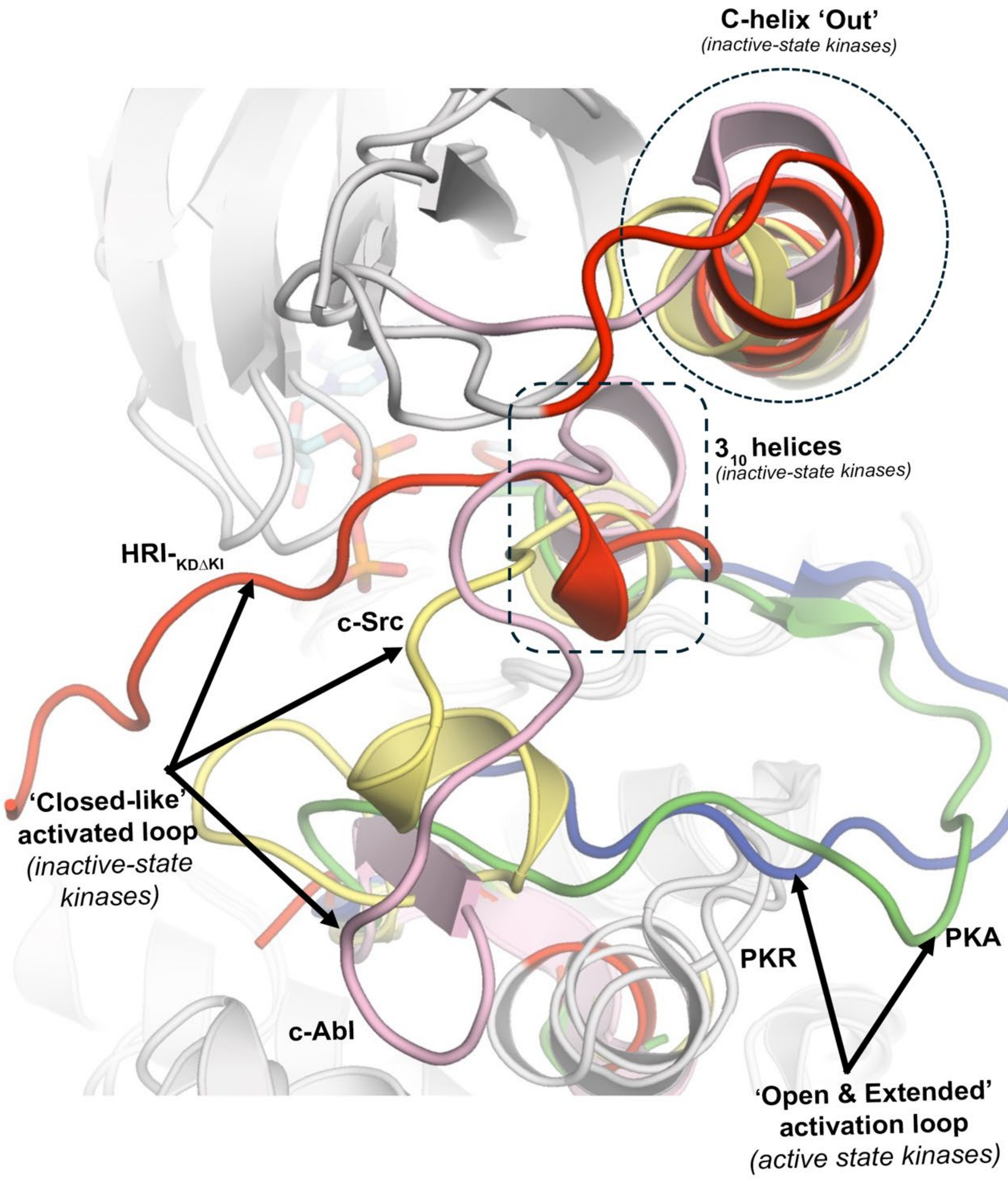

**Figure S10. Structural comparison between $HRI_{KD-\Delta KI}$ and inactive-state conformations of structurally similar kinases.** The structural superimposition between $HRI_{KD-\Delta KI}$ and other similar kinases clearly highlighted both HRI *apo* and MgATP bound crystals structures share an inactive-state configuration as observed in kinases such as c-Src and c-Abl. The key features like C-helix 'Out' (shown in dotted circles), DFG-In conformation as well as 'closed' like conformation for activation loop potentially hindering ATP binding and substrate cleft. The $3_{10}$ helix (shown in dashed rectangles) which arises immediately to C-terminal of DFG motif observed and reported for inactive-state kinase such as c-Src and c-Abl are present in $HRI_{KD-\Delta KI}$ also. For simplicity and consistency, the *apo*- $HRI_{KD-\Delta KI}$ is considered for comparison studies. The open and extended conformation observed in general for the activation loop of active kinases is shown from the PKR (blue cartoon) and PKA kinase (green cartoon).

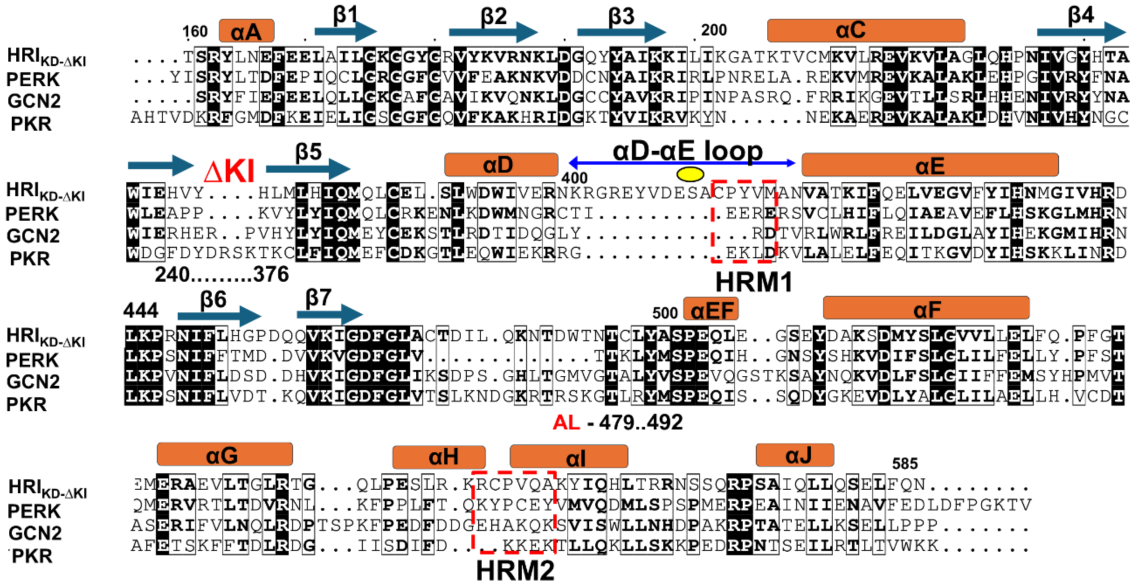

**Figure S11. Sequence-structure based alignment of $HRI_{KD\text{-}\Delta KI}$ with eIF2α kinase family members.** The sequence-structure based comparison studies between kinase domains of $HRI_{KD\text{-}\Delta KI}$ with other eIF2α kinase family members (PERK; *PDB code: 4X7K*), GCN2, *PDB code:7QQ6*, and PKR, *PDB code:2A19*) were carried out using PDBeFold web server. Residues are shaded black where they are identical across all four sequences. The secondary structure elements (helices as orange cylinders and β-strands as blue arrows) are mapped based on the $HRI_{KD\text{-}\Delta KI}$ crystal structure and sequence numbering are according to the $HRI_{KD\text{-}\Delta KI}$ .The location of the deleted KI insert region (ΔKI - 240-376 in red) and disordered activation loop segment (AL - 479-492) is identified by black dotted lines under the sequence alignment. The alignment highlights the sequence context of the αD-αE loop (blue arrow) which is longer than that of related eiF2a homologues is unique to HRI and contains the phosphorylated pSer409 (yellow circle). The position of the two heme regulatory motifs (HRM1 and HRM2) that are unique to $HRI_{KD\text{-}\Delta KI}$ is marked by red dotted-line boxes. The numbering in the alignment is according to HRI (human) sequence (UniprotKB id: Q9BQI3).

### SUPPLEMENTARY TABLES

**Table S2. Data collection and Refinement statistics, Related to Figure 4A**

| | ***HRI$_{KD\text{-}\Delta KI}$*** **PDB Code: 9RHH** | ***HRI$_{KD\text{-}\Delta KI}$ - MgATP*** **PDB Code: (under review, PDB)** |
|---|---|---|
| **Data collection** | | |
| Beamline | Diamond I24 | Diamond I04 |
| Wavelength (Å) | 0.6702 | 0.95374 |
| Resolution range (Å) | 89.000 - 2.08 (2.29 - 2.08) | 58.908 – 2.511 (2.511- 2.715) |
| Space group | P1 21 1 | P1 21 1 |
| a, b, c [ Å] | 56.37, 78.14, 90.17 | 56.85 80.08 88.19 |
| α, β, γ [º] | 90.00, 99.23, 90.00 | 90.00 99.53 90.00 |
| Total reflections[a,b] | 113609 (5827) | 134602 (7028) |
| Unique reflections[a,b] | 32218 (1611) | 20150 (1007) |
| Multiplicity[a,b] | 3.5 (3.6) | 6.7 (7.0) |
| Completeness (%)[a,b] | | |
| *Spherical* | 69.2 (13.6) | 75.3 (18.2) |
| *Ellipsoidal* | 93.1 (68.7) | 90.3 (43.5) |
| $I/\sigma(I)$[a,b] | 8.9 (1.7) | 4.9 (1.4) |
| CC(1/2)[a,b] | 0.997 (0.556) | 0.990 (0.375) |
| $R_{merge}$[a,b] | 0.088 (0.760) | 0.283 (3.217) |
| $R_{pim}$[a,b] | 0.055 (0.467) | 0.118 (1.316) |
| $R_{meas}$[a,b] | 0.104 (0.894) | 0.307 (3.478) |
| **Refinement** | | |
| Resolution Range (Å) | 44.11-2.08 (2.14-2.08) | 58.91-2.511 (2.64 - 2.51) |
| Reflections used in refinement | 32201(105) | 20144 (323) |
| Reflections used for R-free | 1611 (3) | 1016 (21) |
| R-work | 0.206 (.284) | 0.229 (0.298) |
| R-free | 0.232 (.116) | 0.274 (0.301) |
| Protein residues | 538 | 517 |
| Number of atoms | 4373 | 4078 |
| *Macromolecules* | 4129 | 3905 |
| *Ligands* | 27 | 89 |
| *Water* | 217 | 84 |
| Ramachandran Plot(%) | | |
| Favored region | 97.12 | 97.39 |
| Allowed region | 2.69 | 2.61 |
| Outlier region | 0.19 | 0.0 |
| RMS from ideal (bond length) (Å) | 0.002 | 0.002 |
| RMS from ideal (bond angles) (Å) | 0.45 | 0.48 |
| Average B-factor (Å) | 49.53 | 45.66 |
| *Macromolecules* | 49.62 | 45.79 |
| *Solvent* | 46.56 | 35.27 |
| *Ligands* | 60.27 | 49.88 |

[a] These statistics are from the data processed by STARANISO

[b] Values in the parenthesis are for the high-resolution shell
